# Spectral graph theory of brain oscillations

**DOI:** 10.1101/589176

**Authors:** Ashish Raj, Chang Cai, Xihe Xie, Eva Palacios, Julia Owen, Pratik Mukherjee, Srikantan Nagarajan

**Affiliations:** Department of Radiology and Biomedical Imaging, University of California, San Francisco, CA; Department of Bioengineering and Therapeutic Sciences, University of California, San Francisco, CA; Department of Neuroscience, Weill Cornell Graduate School of Medical Sciences, Weill Cornell Medicine, New York, NY; Department of Radiology, University of Washington, Seattle, WA

**Keywords:** spectral graph theory, connectomes, magnetoencephalography, brain activity, alpha rhythm

## Abstract

The relationship between the brain’s structural wiring and the functional patterns of neural activity is of fundamental interest in computational neuroscience. We examine a hierarchical, linear graph spectral model of brain activity at mesoscopic and macroscopic scales. The model formulation yields an elegant closed-form solution for the structure-function problem, specified by the graph spectrum of the structural connectome’s Laplacian, with simple, universal rules of dynamics specified by a minimal set of global parameters. The resulting parsimonious and analytical solution stands in contrast to complex numerical simulations of high dimensional coupled non-linear neural field models. This spectral graph model accurately predicts spatial and spectral features of neural oscillatory activity across the brain and was successful in simultaneously reproducing empirically observed spatial and spectral patterns of alpha-band (8-12 Hz) and beta-band (15-30Hz) activity estimated from source localized scalp magneto-encephalography (MEG). This spectral graph model demonstrates that certain brain oscillations are emergent properties of the graph structure of the structural connectome and provides important insights towards understanding the fundamental relationship between network topology and macroscopic whole-brain dynamics.

**Significance Statement:** The relationship between the brain’s structural wiring and the functional patterns of neural activity is of fundamental interest in computational neuroscience. We examine a hierarchical, linear graph spectral model of brain activity at mesoscopic and macroscopic scales. The model formulation yields an elegant closed-form solution for the structure-function problem, specified by the graph spectrum of the structural connectome’s Laplacian, with simple, universal rules of dynamics specified by a minimal set of global parameters. This spectral graph model demonstrates that certain brain oscillations are emergent properties of the graph structure of the structural connectome and provides important insights towards understanding the fundamental relationship between network topology and macroscopic whole-brain dynamics.

## Introduction

### The Structure-Function Problem in Neuroscience

It is considered paradigmatic in neuroscience that the brain’s structure at various spatial scales is critical for determining its function. In particular, the relationship between the brain’s *structural wiring* and the *functional* patterns of neural activity is of fundamental interest in computational neuroscience. Brain structure and function at the scale of macroscopic networks, i.e. amongst identifiable grey matter (GM) regions and their long-range connections through white matter (WM) fiber bundles, can be adequately measured using current non-invasive measurement techniques. Fiber architecture can be measured from diffusion tensor imaging (DTI) followed by tractography algorithms^1,2^. Similarly, brain function manifested in neural oscillations can be measured non-invasively using magnetoencephalography (MEG) and reconstructed across whole-brain networks. Does the brain’s white matter wiring structure constrain functional activity patterns that arise on the macroscopic network or graph, whose nodes represent gray matter regions, and whose edges have weights given by the structural connectivity (SC) of white matter fibers between them? We address this critical open problem here, as the structural and functional networks estimated at various scales are not trivially predictable from each other^3^.

Although numerical models of single neurons and local microscopic neuronal assemblies, ranging from simple integrate-and-fire neurons to detailed multi-compartment and multi-channel models^4–8^ have been proposed, it is unclear if these models can explain structure-function coupling at meso- or macroscopic scales. At one extreme, the Blue Brain Project^9,10^ seeks to model in detail all 10^11^ neurons and all their connections in the brain. Indeed spiking models linked up via specified synaptic connectivity and spike timing dependent plasticity rules were found to produce regionally and spectrally organized self-sustaining dynamics, as well as wave-like propagation similar to real fMRI data^11^. However, it is unclear whether such efforts will succeed in providing interpretable models at whole-brain scale^12^.

Therefore the traditional computational neuroscience paradigm at the microscopic scale does not easily extend to whole-brain macroscopic phenomena, as large neuronal ensembles exhibit emergent properties that can be unrelated to individual neuronal behavior^13–18^, and are instead largely governed by long-range connectivity^19–22^. At this scale, graph theory involving network statistics can phenomenologically capture structure-function relationships^23–25^, but do not explicitly embody any details about neural physiology^14,15^. Strong correlations between functional and structural connections have also been observed at this scale^3,26–32^, and important graph properties are shared by both SC and functional connectivity (FC) networks, such as small worldness, power-law degree distribution, hierarchy, modularity, and highly connected hubs^24,33^.

A more detailed accounting of the structure-function relationship requires that we move beyond statistical descriptions to mathematical ones, informed by computational models of neural activity. Numerical simulations are available of mean field^17,34,35^ and neural mass^22,36^ approximations of the dynamics of neuronal assemblies. By coupling many such neural field or mass models (NMMs) using anatomic connectivity information, it is possible to generate via large-scale stochastic simulations a rough picture of how the network modulates local activity at the global scale to allow the emergence of coherent functional networks^22^. However, simulations are unable to give an analytical (i.e. closed form) encapsulation of brain dynamics and present an interpretational challenge in that behavior is only deducible indirectly from thousands of trial runs of time-consuming simulations. Consequently, the essential minimal rules of organization and dynamics of the brain remain unknown. Furthermore, due to their nonlinear and stochastic nature, model parameter inference is ill-posed, computationally demanding and manifest with inherent identifiability issues^37^.

How then do stereotyped spatiotemporal patterns emerge from the structural substrate of the brain? How will disease processes perturb brain structure, thereby impacting its function? While stochastic simulations are powerful and useful tools, they provide limited neuroscientific insight, interpretability and predictive power, especially for the practical task of inferring macroscopic functional connectivity from long-range anatomic connectivity. Therefore, there is a need for more direct models of structural network-induced neural activity patterns – a task for which existing numerical modeling approaches, whether for single neurons, local assemblies, coupled neural masses or graph theory, are not ideally suited. Here we use a spectral graph model (SGM) to demonstrate that the spatial distribution of certain brain oscillations are emergent properties of the spectral graph structure of the structural connectome. Therefore, we also explore how the chosen connectome alters the functional activity patterns they sustain.

### A hierarchical, analytic, low-dimensional and linear spectral graph theoretic model of brain oscillations

We present a linear graph model capable of reproducing empirical macroscopic spatial and spectral properties of neural activity. We are interested specifically in the transfer function (defined as the frequency-domain input-output relationship) induced by the macroscopic structural connectome, rather than in the behavior of local neural masses. Therefore we seek an explicit formulation of the frequency spectra induced by the graph, using the eigen-decomposition of the structural graph Laplacian, borrowing heavily from **spectral graph theory** used in diverse contexts including clustering, classification, and machine learning^38–41^. This theory conceptualizes brain oscillations as a linear superposition of eigenmodes. These eigen-relationships arise naturally from a biophysical abstraction of fine-scaled and complex brain activity into a simple linear model of how mutual dynamic influences or perturbations can spread within the underlying structural brain network, a notion that was advocated previously^30,42,43^. We had previously reported that the brain network Laplacian can be decomposed into its constituent “eigenmodes”, which play an important role in both healthy brain function^30,31,44–46^ and pathophysiology of disease^44,47–49^.

We show here that a graph-spectral decomposition is possible at all frequencies, ignoring non-linearities that are operating at the local (node) level. Like previous NMMs, we lump neural populations at each brain region into neural masses, but unlike them we use a linearized (but frequency-rich) local model - see **Figure 1A**. The macroscopic connectome imposes a linear and deterministic modulation of these local signals, which can be captured by a *network transfer function*. The sequestration of local oscillatory dynamics from the macroscopic network in this way enables the characterization of whole brain dynamics deterministically in closed form in Fourier domain, via the eigen-basis expansion of the network Laplacian. As far as we know, this is the first closed-form analytical model of frequency-rich brain activity constrained by the structural connectome.

**Figure 1:**
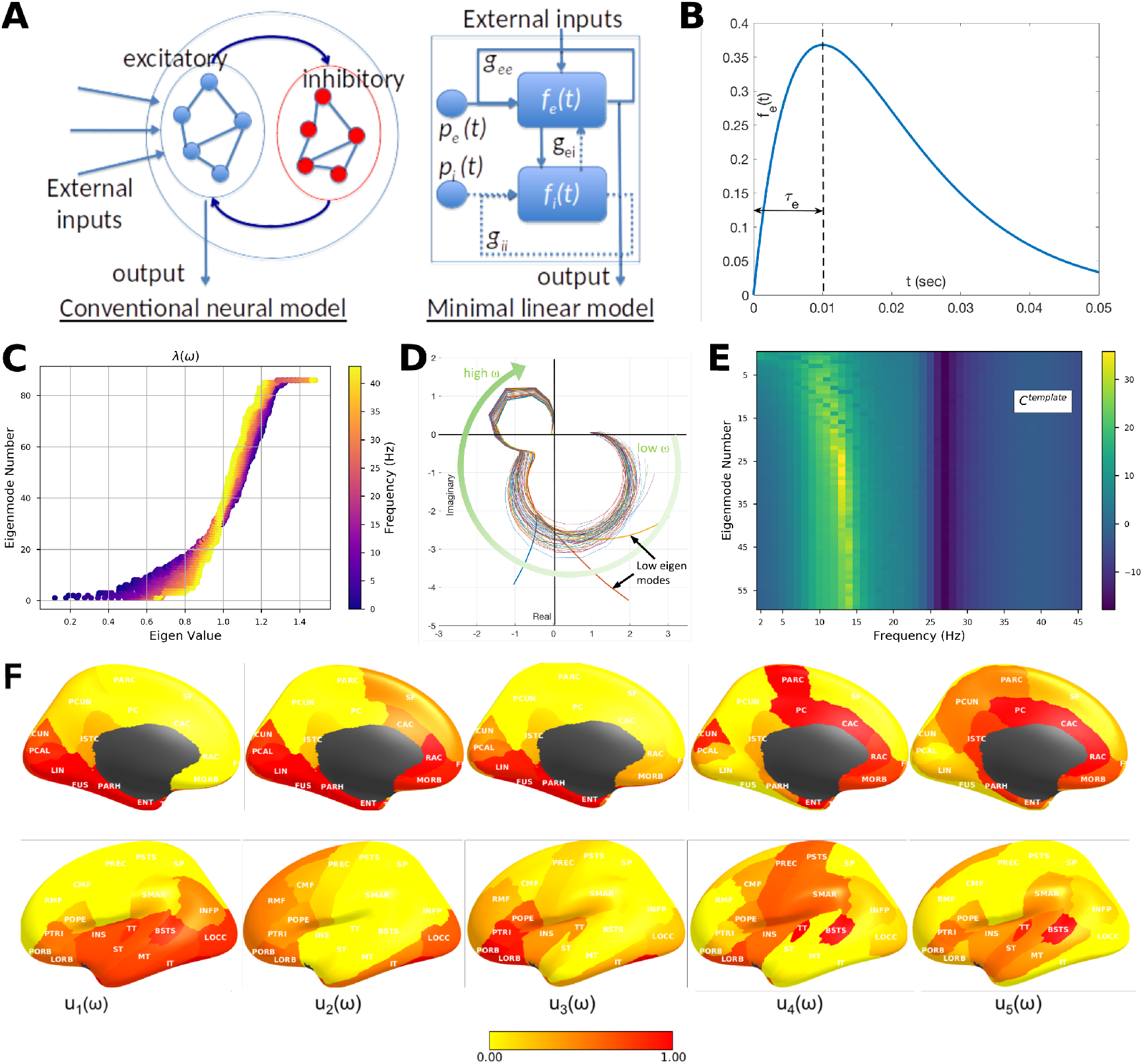
The Linearized Spectral Graph Model. **A:** Conventional neural mass models typically instantiate a large assembly of excitatory and inhibitory neurons, which are modeled as fully connected internally. External inputs and outputs are gated through the excitatory neurons only, and inhibitory neurons are considered strictly local. The proposed linear model condenses these local assemblies into lumped linear systems *f_e_*(*t*) and *f_i_*, Gamma-shaped functions having time constants *τ_e_* and *τ_i_* - see panel **B**. The recurrent architecture of the two pools within a local area is captured by the gain terms *g_ee_*, *g_ii_*, *g_ei_*, indicating the loops created by recurrents within excitatory, inhibitory and cross-populations. **C:** The absolute value of eigenvalues of the complex Laplacian 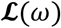 are plotted against the eigenvector index. Each dot represents one eigenvalue *λ*(*ω*); its color represents the frequency *ω* - low (blue) to high (yellow). Clearly, these eigenvalues change somewhat by frequency; small eigenvalues change more compared to large ones. ***D:*** Frequency response of each eigenmode plotted on the complex plane with default choices of model parameters and a template structural connectome from the human connectome project (HCP). Each curve represents the transit in the complex plane of a single eigenmode’s frequency response, starting at low frequencies in the bottom right quadrant, and moving characteristically to the upper left quadrant at high frequencies. The magnitude of the response, given by the distance from the origin, suggests that most eigenmodes have two prominent lobes, roughly corresponding to lower frequency alpha rhythms and higher frequency gamma rhythms, respectively. In contrast, the lowest few eigenmodes start off far from the origin, indicative of a low-pass response. **E:** Magnitude of the frequency response of each eigenmode reinforces these impressions more clearly, with clear alpha peak, as well as slower rhythms of the lowest eigenmodes. The spectral profile of the eigenmodes, especially the peak frequencies, are sensitive to the choice of model parameters. **F:** The spatial patterns of the top 5 eigenmodes of 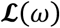, evaluated at the alpha frequency, 10 Hz. The first 4 eigenmodes **u**_1_ – **u**_4_, produce posterior and temporal spatial patterns, including many elements of the **default mode network; u**_4_ resembles the **sensorimotor network**; and **u**_5_ the **structural core** of the human connectome. However, these patterns are not exclusive and greatly depend on the frequency at which they are evaluated, as well as the model parameters and the connectome.

We applied this model to and validated its construct against measured source-reconstructed MEG recordings in healthy subjects under rest and eyes-closed. The model closely matches empirical spatial and spectral MEG patterns. In particular, the model displays prominent alpha and beta peaks, and, intriguingly, the eigenmodes corresponding to the alpha oscillations have the same posterior-dominant spatial distribution that is repeatedly seen in eyes-closed alpha power distributions. In contrast to existing less parsimonious models in the literature that invoke spatially-varying parameters or local rhythm generators, to our knowledge, this is the first account of how the spectral graph structure of the structural connectome can parsimoniously explain the spatial power distribution of alpha and beta frequencies over the entire brain measurable on MEG.

## Methods

### Spectral graph model development

#### Notation

In our notation, vectors and matrices are represented by boldface, and scalars by normal font. We denote frequency of a signal, in Hertz, by symbol *f*, and the corresponding angular frequency as *ω* = 2*πf*. The connectivity matrix is denoted by ***C*** = {*c_jk_*}, consisting of connectivity strength *c_ij_* between any two pair of regions *j*, *k*.

#### Model summary

Details of the Spectral Graph model (SGM) is described in detail below. There are very few model parameters, seven in total: *τ_e_*, *τ_i_*, *τ_G_*, *v*, *g_ii_*, *g_ei_*, *α*, which are all global and apply at every node. See **Table 1** for their meaning, initial value and range. Note that the entire model is based on a single equation of graph dynamics, Eq (1), which is repeatedly applied to each level of the hierarchy. Here we used two levels: a mesoscopic level where connectivity is all-to-all, and a macroscopic level, where connectivity is measured from fiber architecture. In theory, this template could be refined into finer levels, where neural responses become increasingly non-linear, and connectivity becomes sparser and structured.

**Table 1:** SGM parameters values and limits.

| Name | Symbol | Initial/default Value | Lower/Upper bound for optimization |
| --- | --- | --- | --- |
| Excitatory Time constant | $\tau_e$ | 12 ms | [5ms, 20ms] |
| Inhibitory Time constant | $\tau_i$ | 3 ms | [5ms, 20ms] |
| Graph Time constant | $\tau_G$ | 6 ms | [5ms, 20ms] |
| Excitatory gain | $g_{ee}$ | 1 | n/a |
| Inhibitory gain | $g_{ii}$ | 1 | [0.5, 5] |
| Excitatory gain | $g_{ei}$ | 4 | [0.5, 5] |
| Transmission velocity | $v$ | 5 m/s | [5 m/s, 20 m/s] |
| Long-range connectivity coupling constant | $\alpha$ | 1 | [0.1, 1] |

#### Canonical rate model over a graph

We use a canonical rate model to describe neural activity across two hierarchical levels - local cortical mesoscopic levels and long-range macroscopic levels. At each level of the hierarchy of brain circuits, we hypothesize a simple linear rate model of recurrent reverberatory activity given by

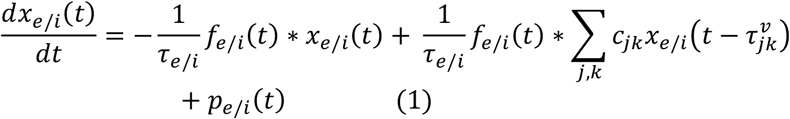

where *x*_*e*/*i*_(*t*) is the mean signal of the excitatory/inhibitory populations and *p*_*e*/*i*_(*t*) is internal noise source reflecting local cortical column computations or input. The transit of signals, from pre-synaptic membranes, through dendritic arbors and axonal projections, is sought to be captured into ensemble average neural impulse response functions 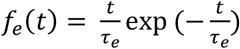 and 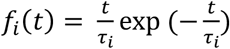 respectively. We disregard the non-linearity of the neural response, hence the output at the terminal to a presynaptic input *u*(*t*) is the simple convolution *x_e_*(*t*) = *f_e_*(*t*) * *u*(*t*). The neural responses *f*_*e*/*i*_(*t*) are Gamma-shaped responses (**Figure 1B**) parameterized by time constants *τ*_*e*/*i*_ that here represent the end result of both synaptic membrane capacitance and the distribution of dendritic/axonal delays introduced by the arborization. NMMs typically use a single classical exponential decay term for membrane capacitance only, since NMMs model highly local cell assemblies where multisynaptic connections are infrequent and axonal and dendritic transport delays are usually incorporated explicitly via connectivity weights and delays. Since our lumped model was designed for relatively large cortical regions, we employ the Gamma-shaped *f*_*e*/*i*_ to capture not just classical membrane capacitance but also the expected diversity of dendritic transport delays. The dynamics of the entire assembly modeled via a self-decaying term 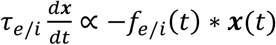, typically used in most rate or NMM models, but the difference here is that we chose to apply convolution with neural response *f*_*e*/*i*_(*t*) within the decay process. We believe this is necessary to ensure that the dynamics of the population cannot participate in the internal recurrent dynamics of the region until the signal has passed through one instance of the neuronal response. Since this neural response is meant to capture a distribution of local circuit delays, its time constants *τ*_*e*/*i*_ are purposefully far longer (up to 20ms) than expected from membrane capacitance alone. Studies of cortical lag times using paired electrode recordings between primary and higher cortices demonstrate this. A short visual stimulus causes a neural response in the ferret V1 within 20ms post-stimulus, in the primary barrel field within 16-36ms, and the entire visual cortex becomes engaged 48-70ms after stimulus^6^. Brief deflection of a single barrel whisker in the mouse evokes a somatotopically mapped cortical depolarization that remains localized to its C2 barrel column only for a few milliseconds, thence rapidly spreading to a large part of sensorimotor cortex within tens of milliseconds, a mechanism considered essential for the integration of sensory information^50,51^. Interestingly, the evoked response curve in S1 from the^50^ study had a prominent Gamma shape. Of note, the duration of S1 response (~50ms) was considerably longer than the time to first sensory response in C2 (7.2ms)^50^. Interestingly, feedback projections from higher to lower areas take ~50ms, hence have a much slower apparent propagation velocity (0.15-0.25m/s) than what would be predicted by axonal conduction alone (1-3m/s)^6^.

Individual neural elements are connected to each other via connection strengths *c_jk_*. Let the cortico-cortical fiber conduction speed be *v*, which here is assumed to be a global constant independent of the pathway under question. For a given pathway connecting regions *j* and *k* of length *d_jk_*, the conduction delay of a signal propagating from region *j* to region *k* will be given by 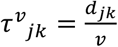. Hence signals from neighboring elements also participate in the same recurrent dynamics, giving the 2^nd^ term of Eq (1). Equation (1) will serve as our canonical rate model, and will be reproduced at all levels of the hierarchy, and only the connectivity strengths will vary depending on the level of hierarchy we are modeling, as explained below.

#### Local neural assemblies

The local connectivities 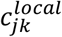 are assumed to be all-to-all, giving a complete graph. Further, the axonal delays 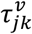 associated with purely local connections were already incorporated in the lumped impulse responses *f*_*e*/*i*_(*t*). Hence, we assert:

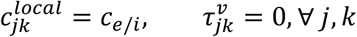

From spectral graph theory, a complete graph has all equal eigenvalues which allows the local network to be lumped into gain constants, and the summation removed. Indeed, rewriting *x*_*e*/*i*_(*t*) as the mean signal of all the excitatory/inhibitory cells and setting the gains *g_ee_* = 1 − *c_e_N_e_* and *g_ii_* = 1 − *c_i_*/*N_i_* we get

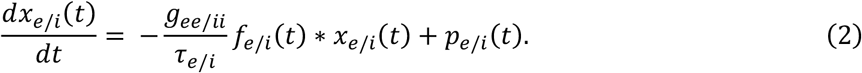

Given the Fourier Transform pairs 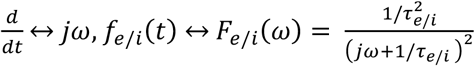, we take the Fourier transform of Eq(1) and obtain the local assembly’s frequency spectrum:

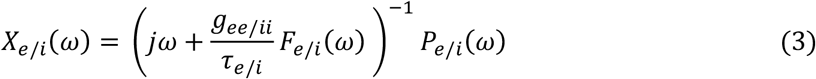

Writing this in terms of transfer functions *X_e_*(*ω*) = *H_e_*(*ω*)*P_e_*(*ω*), *X_i_*(*ω*) = *H_i_*(*ω*)*P_i_*(*ω*) we get the lumped local system illustrated in **Figure 1A**. Finally, we must also account for signals that alternate between the two populations, which is given by the transfer function

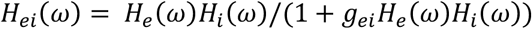

We fix *g_ee_* = 1 without loss of generality, and let the other terms *g_ii_*, *g_ei_* be model parameters to be fitted. Finally, the total cortical transfer function is the sum

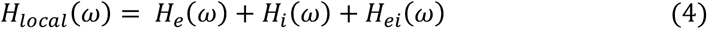

and *X_local_*(*ω*) = *H_local_*(*ω*)*P*(*ω*) represents all neural activity in this region, whether from excitatory or inhibitory cells. The canonical local activity is therefore defined by the Fourier transform pair: *X_loca1_*(*t*) ↔ *X_local_*(*ω*)

#### Macroscopic scale: signal evolution on the entire graph

For the macroscopic level, we use the same canonical network dynamics as Eq (1), but now the inter-regional connectivity *c_jk_* is non-zero and given by the structural connectome. Similarly, axonal conductance delays are determined by fiber length and conductance speed 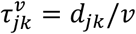. Further, the external driving signals at each node is the local neural activity *x_local_*(*t*) defined above rather than a noise process *p*(*t*). In the interest of parsimony we set each node of the macroscopic graph to have the same internal power spectrum *X_local_*(*ω*) - i.e. all regions are experiencing the same transfer function, driven by identically distributed (but of course not identical) noise. At this scale, activity measured at graph nodes is no longer excitatory or inhibitory, but mixed, and the corticocortical connections are all between long, pyramidal excitatory-only cells. Thus, for the k-th node

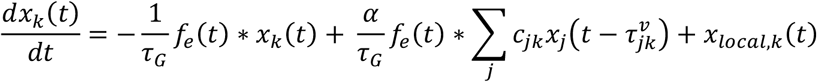

Here we have introduced a global coupling constant *α*, similar to most connectivity-coupled neural mass models, that seeks to control the relative weight given to long-range afferents compared to local signals. We have also introduced a new time constant, *τ_G_*, which is an excitatory time constant and it may be the same as the previously used constant *τ_e_*. However, we allow the possibility that the long-range projection neurons might display a different capacitance and morphology compared to local circuits, hence we have introduced *τ_G_* (subscript G is for “graph” or “global”).

Stacking all equations from all nodes and using vector valued signals ***x***(*t*) = {*x_k_*(*t*)}, we can write

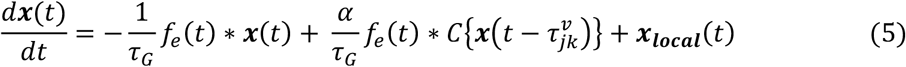

where the braces {·} represent all elements of a matrix indexed by *j*, *k*.

We wish to evaluate the frequency spectrum of the above. In Fourier space, delays become phases; hence we use the transform pairs 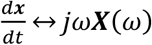 and ***x***(*t* − *τ*) ↔ *e*^−*jτω*^ ***X***(*ω*). Therefore, define a “complex connectivity matrix” at any given angular frequency *ω* as ***C***\*(*ω*) = {*c_jk_* exp(−*jω τ^v^_jk_*)}. We then define a normalized complex connectivity matrix at frequency *ω* as

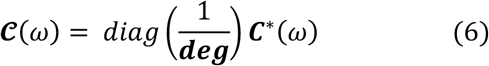

where the degree vector ***deg*** is defined as *deg_k_* = ∑_*j*_ *c_jk_*. Taking the Fourier transform of Eq (5), we get

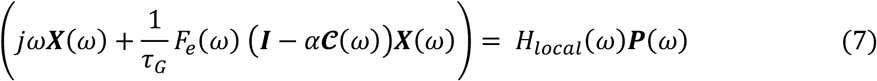

where we assumed identically distributed noise signals driving both the excitatory and inhibitory local populations at each node, such that *P_e,k_*(*ω*) = *P_i,k_*(*ω*) = *P_k_*(*ω*) at the k-th node. We then collected all nodes’ driving inputs in the vector ***P***(*ω*) = {*P_k_*(*ω*), ∀*k*}. Here, we define the complex Laplacian matrix

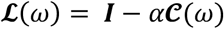

where ***I*** is the identity matrix of size *N* × *N*. This complex Laplacian will be evaluated via the eigen-decomposition

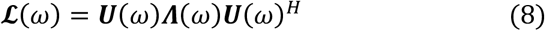

where ***Λ***(*ω*) = *diag*([*λ*_1_(*ω*), …, *λ_N_*(*ω*)]) is a diagonal matrix consisting of the eigenvalues of the complex Laplacian matrix of the connectivity graph 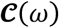, at the angular frequency *ω*.

Hence

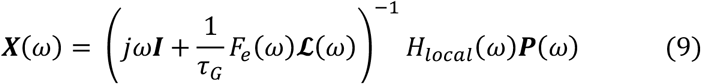

where we invoke the eigen-decomposition of 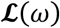, and that ***U***(*ω*)***U***(*ω*)^***H***^ = ***I***. It can then be shown easily that

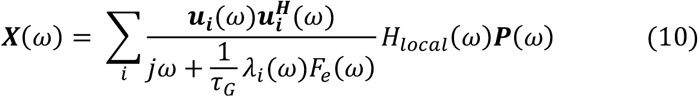

This is the steady state frequency response of the whole brain dynamics. In steady state, we assume that each cortical region is driven by internal noise that spans all frequencies, i.e. white noise. Hence, we assume that the driving function ***p***(*t*) is an uncorrelared Gaussian noise process, such that 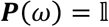, where 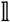 is a vector of ones. This asserts identical cortical responses at each brain region.

### Experimental Procedures

#### Study cohort

We acquired MEG, anatomical MRI, and diffusion MRI for 36 healthy adult subjects (23 males, 13 females; 26 left-handed, 10 right-handed; mean age 21.75 years (range: 7-51 years). All study procedures were approved by the institutional review board at the University of California at San Francisco (UCSF) and are in accordance with the ethics standards of the Helsinki Declaration of 1975 as revised in 2008.

#### MRI

A 3 Tesla TIM Trio MR scanner (Siemens, Erlangen, Germany) was used to perform MRI using a 32-channel phased-array radiofrequency head coil. High-resolution MRI of each subject’s brain was collected using an axial 3D magnetization prepared rapid-acquisition gradient-echo (MPRAGE) T1-weighted sequence (echo time [TE] = 1.64 ms, repetition time [TR] = 2530 ms, TI = 1200 ms, flip angle of 7 degrees) with a 256-mm field of view (FOV), and 160 1.0-mm contiguous partitions at a 256×256 matrix. Whole-brain diffusion weighted images were collected at b = 1000*s*/*mm*^2^ with 30 directions using 2mm voxel resolution in-plane and through-plane.

#### Magneto-encephalography (MEG) data

MEG recordings were acquired at UCSF using a 275-channel CTF Omega 2000 whole-head MEG system from VSM MedTech (Coquitlam, BC, Canada). All subjects were instructed to keep their eyes closed for five minutes while their MEGs were recorded at a sampling frequency of 1200 Hz.

### Data Processing

#### Region Parcellations

The T1-weighted images were parcellated into 68 cortical regions and 18 subcortical regions using the using the Desikan-Killiany atlas available in the FreeSurfer software^52^. To do this, the subject specific T1-weighted images were back-projected to the atlas using affine registration, as described in our previous studies^18,53^.

#### Structural Connectivity Networks

We constructed different structural connectivity networks with the same Desikan-Killiany parcellations to access the capabilities of our proposed model. Firstly, we obtained openly available diffusion MRI data from the MGH-USC Human Connectome Project to create an average template connectome. As in our previous studies^18,53^, subject specific structural connectivity was computed on diffusion MRI data: *Bedpostx* was used to determine the orientation of brain fibers in conjunction with *FLIRT*, as implemented in the *FSL* software^54^. In order to determine the elements of the adjacency matrix, we performed tractography using *probtrackx2*. We initiated 4000 streamlines from each seed voxel corresponding to a cortical or subcortical gray matter structure and tracked how many of these streamlines reached a target gray matter structure. The weighted connection between the two structures *c_i,j_*, was defined as the number of streamlines initiated by voxels in region *i* that reach any voxel within region *j*, normalized by the sum of the source and target region volumes 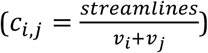. This normalization prevents large brain regions from having high connectivity simply due to having initiated or received many streamlines. Afterwards, connection strengths are averaged between both directions (*c_i,j_* and *c_j,i_*) to form undirected edges. It is common in neuroimaging literature to threshold connectivity to remove weakly connected edges, as this can greatly influence the implied topology of the graph. In our work, we chose not to apply further thresholding, as unlike conventional graph theoretic metrics, linear models of spread and consequently network eigenmodes are relatively insensitive to implied topology induced by presence (or lack) of weak nonzero connections. However, to determine the geographic location of an edge, the top 95% of non-zero voxels by streamline count were computed for both edge directions. The consensus edge was defined as the union between both post-threshold sets.

#### MEG processing and source reconstruction

MEG recordings were down-sampled from 1200 Hz to 600 Hz, then digitally filtered to remove DC offset and any other noisy artifact outside of the 1 to 160 Hz bandpass range. Since MEG data are in sensor space, meaning they represent the signal observable from sensors placed outside the head, this data needs to be “inverted” in order to infer the neuronal activity that has generated the observed signal by solving the so-called inverse problem. Several effective methods exist for performing *source localization*^55–57^. Here we eschew the common technique of solving for a small number of discrete dipole sources which is not fully appropriate in the context of inferring resting state activity, since the latter is neither spatially sparse not localized. Instead, we used adaptive spatial filtering algorithms from the NUTMEG software tool written in house^58^ in MATLAB (The MathWorks, Inc., Natick, Massachusetts, United States). To prepare for source localization, all MEG sensor locations were co-registered to each subject’s anatomical MRI scans. The lead field (forward model) for each subject was calculated in NUTMEG using a multiple local-spheres head model (three-orientation lead field) and an 8 mm voxel grid which generated more than 5000 dipole sources, all sources were normalized to have a norm of 1. Finally, the MEG recordings were projected into source space using a beamformer spatial filter. Source estimates tend to have a bias towards superficial currents and the estimates are more error-prone when we approach subcortical regions, therefore, only the sources belonging to the 68 cortical regions were selected to be averaged around the centroid. Specifically, all dipole sources were labeled based on the Desikan-Killiany parcellations, then sources within a 20 mm radial distance to the centroid of each brain region were extracted, the average time course of each region’s extracted sources served as empirical resting-state data for our proposed model.

#### Alternative benchmark model for comparison

In order to put the proposed model in context, we also implemented for comparison a Wilson-Cowan neural mass model^17,35,37,59^ with similar dimensionality. Although NMMs like this can and have been implemented with regionally varying local parameters, here we enforced uniform, regionally non-varying local parameters, meaning all parcellated brain regions shared the same local and global parameters. This is a fair comparison since the proposed model is also regionally non-varying. The purpose of this exercise is to ascertain whether a non-regional NMM can also predict spatial power variations purely as a consequence of network transmission, like the proposed model, using the same model optimization procedure (see below). This NMM incorporates a transmission velocity parameter that introduces a delay based on fiber tract lengths extracted from diffusion MRI, but, unlike our model, does not seek to explicitly evaluate a frequency response based on these delays.

### Model Optimization

We computed *maximum a posteriori* estimates for parameters under a flat non-informative prior. A simulated annealing optimization algorithm was used for estimation and provided a set of optimized parameters {*τ_e_*, *τ_i_*, *τ_c_*, *g_ei_*, *g_ii_*, *α*, *v*}. We defined a data likelihood or goodness of fit (GOF) as the Pearson correlation between empirical source localized MEG power spectra and simulated model power spectra, averaged over all 68 regions of a subject’s brain. The proposed model has only seven global parameters as compared to neural mass models with hundreds of parameters, and is available in closed-form. To improve the odds that we capture the global minimum, we chose to implement a probabilistic approach of simulated annealing^60^. The algorithm samples a set of parameters within a set of boundaries by generating an initial trial solution and choosing the next solution from the current point by a probability distribution with a scale depending on the current “temperature” parameter. While the algorithm always accepts new trial points that map to cost-function values lower than the previous cost-function evaluations, it will also accept solutions that have cost-function evaluations greater than the previous one to move out of local minima. The acceptance probability function is 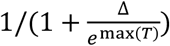, where T is the current temperature and Δ is the difference of the new minus old cost-function evaluations. The initial parameter values and boundary constraints for each parameter are given in Supplementary Table 1. All simulated annealing runs were allowed to iterate over the parameter space for a maximum of *N_p_* × 3000 iterations, where *N_p_* is the number of parameters in the model. As a comparison, we performed the same optimization procedure to a regionally non-varying Wilson-Cowan neural mass model^35,59^. We have recently reported a similar simulated annealing optimization procedure on this model^37^.

## Results

### Graph Laplacian eigenmodes mediate a diversity of frequency responses

First, we demonstrate the spectra produced by graph eigenmodes as per our theory using default choices of model parameters. **Figure 1C** shows the eigen-spectrum of the complex Laplacian, with eigenvalue magnitude ranging from 0 to 1. The absolute value of eigenvalues of the complex Laplacian 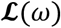 are plotted against the eigenvector index. Each dot represents one eigenvalue *λ*(*ω*); its color represents the frequency *ω* - low (blue) to high (yellow). Clearly, these eigenvalues change somewhat by frequency. Small eigenvalues undergo a larger shift due to frequency, while the large ones stay more stable and tightly clustered around the nominal eigenvalue (i.e. at *ω* = 0). Each eigenmode produces a frequency response based on its frequency-dependent eigenvalue (**Figure 1D, E**). **Figure 1D** shows the transit in the complex plane of a single eigenmode’s frequency response, starting at low frequencies in the bottom right quadrant, and moving to the upper left quadrant at high frequencies. The magnitude, given by distance from origin, suggests that most eigenmodes have two prominent lobes, one roughly corresponding to lower frequency alpha rhythm and another corresponding to higher frequency beta or gamma rhythms, respectively. In contrast, the lowest few eigenmodes start off far from the origin, indicative of a low-pass response. The magnitude of these complex-valued curves shown in figure 1E reinforces these impressions, with clear alpha peak, as well as slower rhythms of the lowest eigenmodes. The spectral profile of the eigenmodes, especially the peak frequencies, are sensitive to the choice of model parameters as demonstrated below.

The spatial patterns of the first 5 eigenmodes of 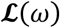, evaluated at the alpha peak of 10 Hz, are shown in **Figure 1F**. Eigenmodes **u_1−4_** produce posterior and temporal spatial patterns, including many elements of the **default mode network; u**_4_ resembles the **sensorimotor network**; and **u**_5_ the **structural core** of the human connectome. However, these patterns are not exclusive and greatly depend on the frequency at which they are evaluated, as well as the model parameters. Higher eigenmodes are especially sensitive to axonal velocity and frequency (not shown here).

Since the spectral graph model (SGM) relies on connectome topology, we demonstrate in **Figure 2** that different connectivity matrices produce different frequency responses: A) the individual’s structural connectivity matrix, B) HCP average template connectivity matrix, C) uniform connectivity matrix of ones, D) a randomly generated matrix, E) and F) are randomly generated matrices with 75% and 95% sparsity respectively. For figure 2A, optimized parameters for the individual subject’s connectome were used. For figures 2B-F, parameters optimized for the HCP template were used. We can observe the spectral profile of the eigenmodes, especially the peak frequencies, are sensitive to the choice of the connectome and the model parameters. All modeled power spectra show a broad alpha peak at around 10 Hz and a narrower beta peak at around 20 Hz. This is expected, since these general spectral properties are governed by the local linearized neural mass model. It is important to note that different eigenmodes accommodate a diversity of frequency responses; for instance, the lowest eigenmodes show a low-frequency response with no alpha peak whatsoever. In the frequency responses from biologically realistic individual and HCP template connectomes, there is a diversity of spectral responses amongst eigenmodes that is lacking in the response produced by the unrealistic uniform and randomized matrices. As we will see below, graph topology is critical to the power spectrum it induces, hence we explored whether and how sparsity of random graphs mediates spectral power (**Figure 2D-F**). At incrementally increasing sparsity levels, the diversity of spectral responses of different eigenmodes increases and approaches that of realistic connectomes. Therefore, graph eigenmodes induce unique and diverse frequency responses that depend on the topology of the graph.

**Figure 2:**
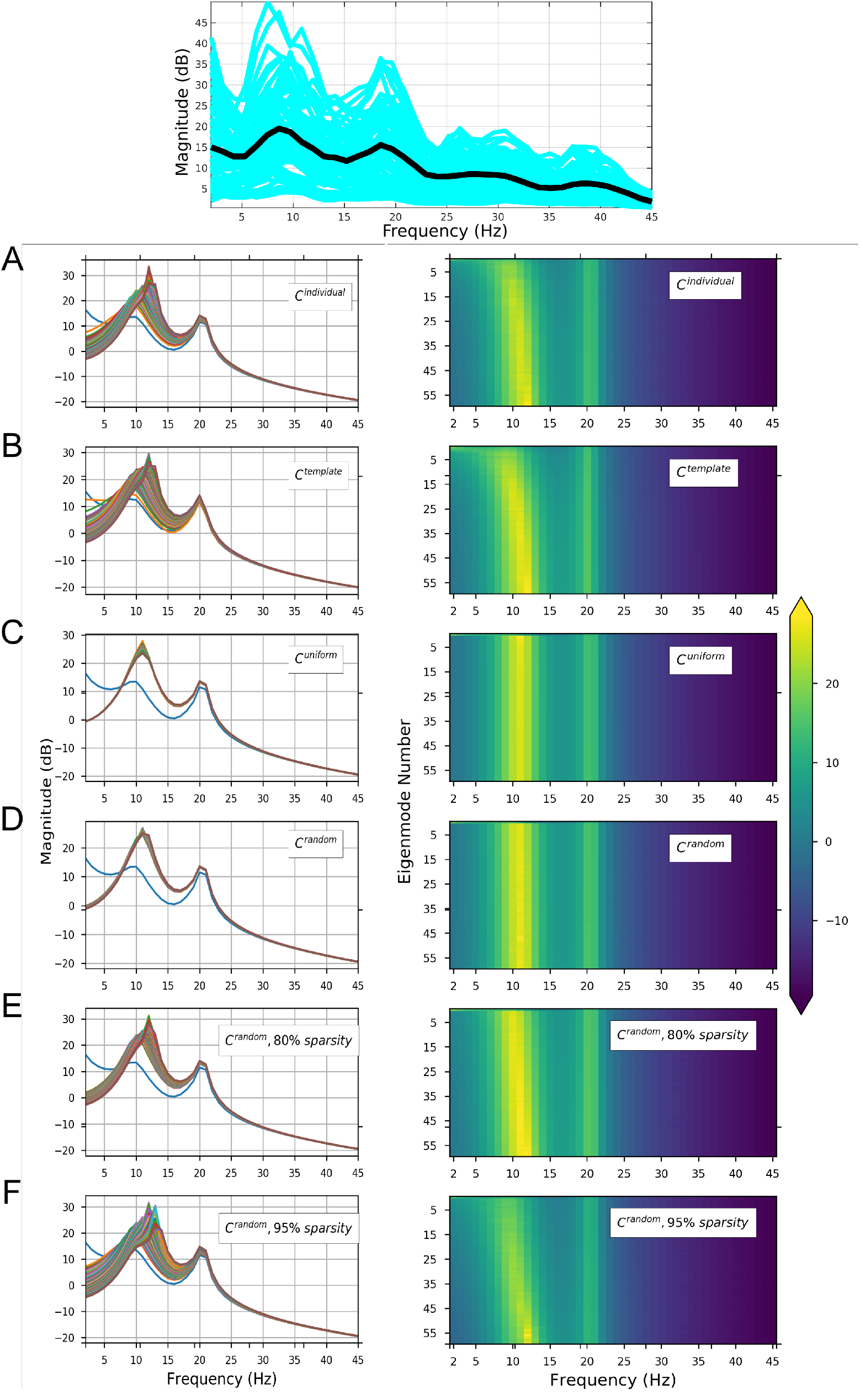
Spectral graph model predictions of MEG spectra for one representative subject. Top - Observed MEG power spectrum for each of the 68 parcellated brain regions. Average spectra for each brain region are shown in blue, and the average spectrum across all brain regions is shown in thick black curve. The subsequent rows show each eigenmode’s spectral magnitude response with model parameters optimized to match the observed spectrum (*τ_e_* = 0.0073, *τ_i_* = 0.0085, *τ_G_* = 0.0061, *g_ei_* = 2.9469 *g_ii_* = 4.4865, *v* = 18.3071 and *α*. = 0.4639). Left column shows each eigenmode’s frequency response in a differently-colored curve, while the right column shows the same information as a heatmap. **A:** Model using subject’s individual structural connectivity matrix. **B:** Model using a template structural connectivity matrix obtained by averaging structural connectivity from 80 HCP subjects. **C:** Model using uniform connectivity matrix of ones. **D:** Model using randomized connectivity matrix with no sparsity. **E:** Model using randomized connectivity matrix with 75% sparsity. **F:** Model using randomized connectivity matrix with 95% sparsity. In all cases the connectome modulates the spectral response in delta-beta range, leaving the higher gamma frequencies unchanged. In general, the low eigenmodes (**u_1_** – **u_20_**) appear to modulate the lower frequency range, up to beta, and may be considered responsible for the diversity of spectra observed in the model.

### Spectral distribution of MEG power depends on model parameters but not connectivity

Network eigenmodes exhibit strong spatial patterning in their frequency responses, even with identical model parameters (**Figure 3**). We evaluated the model spectral response using the subject-specific *C^ind^* matrices of 4 representative subjects (**Figure 3A**). The model power spectra strikingly resemble empirical MEG spectra, displaying both the alpha and beta peaks on average, and similar regional variability as in real data.

Regional averages of empirical and modeled power spectra of the entire group after full parameter optimization over individual subjects are shown in **Figure 3B**. The model closely replicates the observed power spectrum (red circles) equally well with both *C^ind^*(black triangles) and *C^HCP^* (purple triangles). Thus, in most cases we can safely replace the subject-specific connectome with the template connectome. In contrast, when non-optimized average parameters were used (golden green triangles), it resulted in a worse fit, especially at high frequencies, suggesting that individualized parameter optimization is essential to produce realistic spectra. We also examined the model behavior for a random connectome (bright green triangles) or a distance-based connectome (blue triangles) was chosen with identical sparsity (80%) to the actual connectome, and found that even with optimized parameters the average spectra could be accounted for by these connectomes.

**Figure 3:**
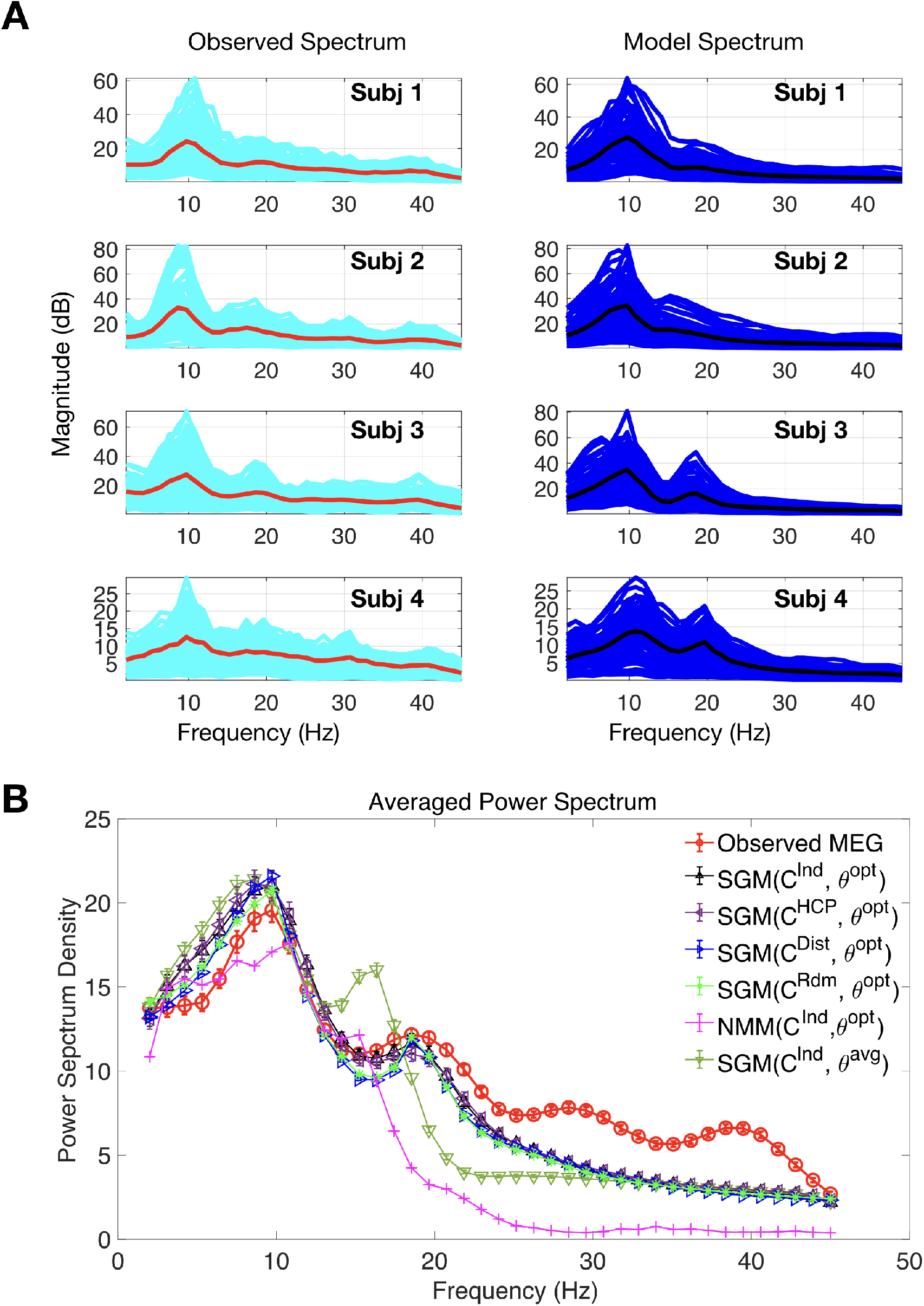
Spectral graph model depicts MEG spectra across subjects. **A:** The observed spectra and spectral graph model’s simulated spectra for four representative subjects. Red and cyan curves illustrate source localized empirical average spectra and region-wise spectra respectively, while black and blue curves illustrate modeled average spectra and region-wise spectra respectively. **B:** Averaged observed spectrum across subjects is shown in red. The average simulated model spectra summing the first two-third eigenmodes with optimized parameters for individual subject’s connectome is shown in black. Model spectrum with optimized parameters and the HCP template connectome is shown in purple. Model spectrum with average parameter values and individual subject’s connectome is shown in golden green. Model spectrum with optimized parameters and a distance connectome with 80% sparsity is shown in blue. Model spectrum with optimized parameters and symmetric random connectomes with 80% sparsity is shown in green. Finally, model power spectrum estimated by a neural mass model (NMM) with each subject’s optimized global parameters and a HCP template connectome is shown in pink.

As another benchmark for comparison, a non-linear neural mass model^35,59^ using our in-house MATLAB implementation^37^, was generally able to produce characteristic alpha and beta frequency peaks, but this model does not resemble empirical wideband spectra. Note that no regionally-varying NMM parameters were used in order to achieve a proper comparison with our model, but both models were optimized with the same algorithm.

**Figure 4A** shows violin plots of the optimized values, indicating that there is a large range of individually optimal model parameters across subjects. The time constants *τ_e_*, *τ_i_* showed tight clustering but the rest of the parameters showed high variability across subjects. The optimal parameters are in a biologically plausible range, similar to values reported in numerous neural mass models. The optimization algorithm aimed to maximize a cost function proportional to the posterior likelihood of the model, and was quantified by the Pearson’s correlation between MEG and modeled spectra (“Spectral correlation”). The convergence plots shown in **Figure 4B**, one curve for each subject, indicates substantial improvement in cost function from default choice as optimization proceeds. The distribution of optimized spectral correlations is shown in **4C**. Other model choices were evaluated for comparison: SGM on random connectomes with and without a distance effect described in Methods, and SGM applied with average optimal model parameters instead of individually optimized ones. In order to test for significance, Fisher’s R to z transform was applied, followed by a paired t-test across all subjects between the optimal SGM and other models. The spectral fits for an SGM model with individual connectomes were significantly better than SGM models with average parameters, no matter what connectomes were chosen (p<0.001). Interestingly, spectral fits for SGM model were comparable across all connectomes (p>0.05). Furthermore, spectral fits for the SGM model were significantly better than that for NMM models with optimized parameters and individual connectome (p<*). Therefore, we conclude that with the graph spectral model, the overall regional spectra appear to be dependent on global model parameters rather than on the actual structural connectome.

**Figure 4:**
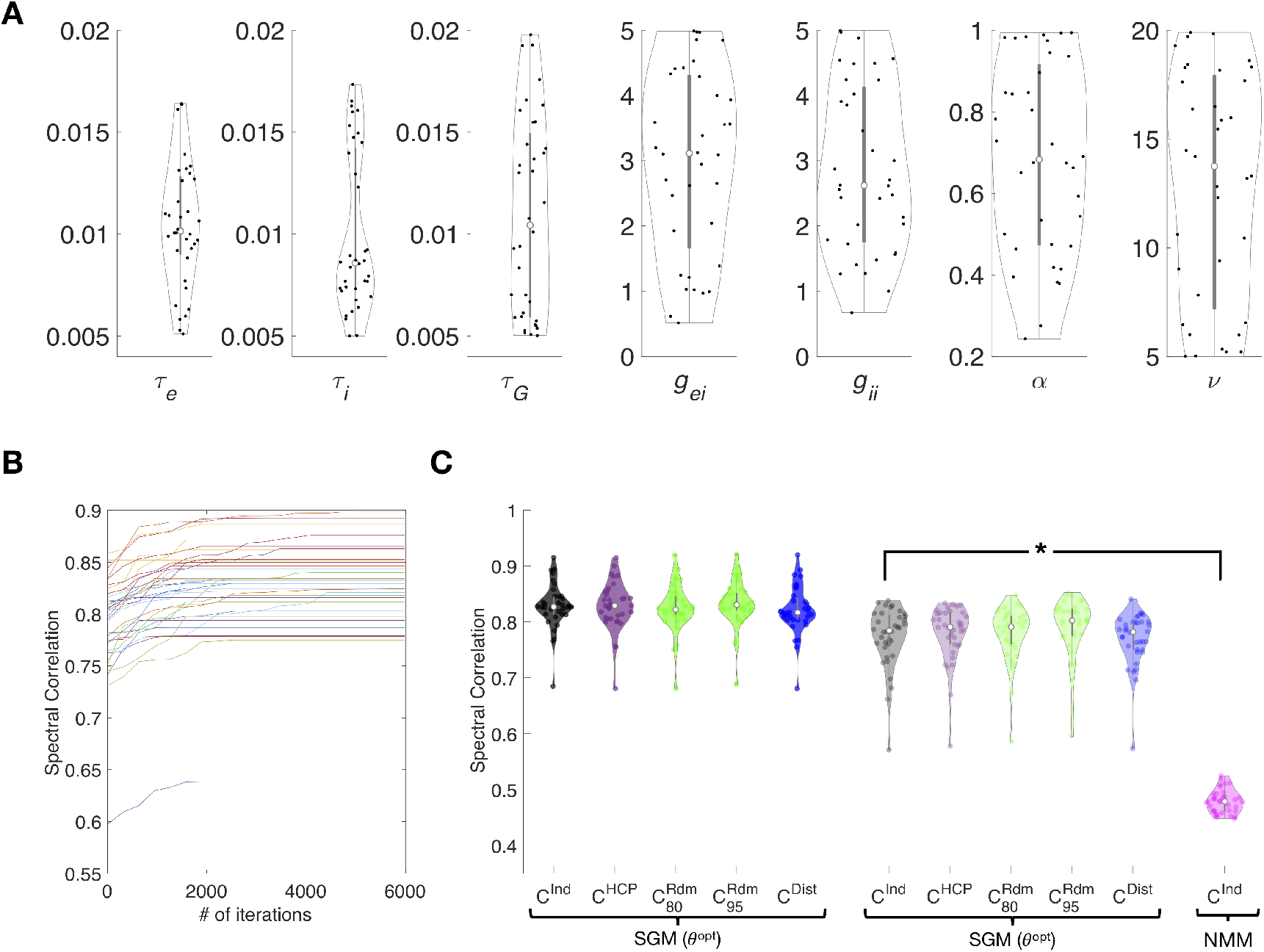
Spectral graph model parameter optimization improves spectral fits. **A:** Distribution of optimized model parameter values across all 36 subjects for the set of parameters {*τ_e_*, *τ_i_*, *τ_c_*, *g_ei_*, *g_ii_*, *α*, *v*} are shown in violin plots with each dot representing one subject. **B.** Performance of optimization algorithm. Spectral Pearson correlation between model and source localized MEG spectra at each iteration. Each curve shows the spectral correlation achieved by the model optimized for a single subject, averaged over all regions, with increasing mean correlation values until the algorithm convergence to a set of optimized parameters. **C:** Distribution of spectral correlations between optimized model and observed spectra across subjects. Correlations with optimized parameters are shown in the left three columns with individual connectomes (black), symmetric random connectomes with 80% and 95% sparsity (blue) and geodesic distance-based connectomes (green). Correlation with average parameter values and individual connectomes are shown in golden green. Spectral correlations are highest for the SGM model with optimized parameters and the individual subject specific connectome when compared to SGM model with average parameters, regardless of the connectome and with an optimized NMM model, as denoted by asterisk (p<0.001).

### Spectral graph model recapitulates the spatial distribution of MEG power

Next, we establish that the model is able to reproduce region-specific spectra, even though it uses identical local oscillations. We integrated the spectral area in the range 8-12 Hz for alpha and 13-25 Hz for beta, of each brain region separately. We define “*spatial correlation*” (as compared to spectral correlation above) as Pearson’s R between the *regional distribution* of empirical MEG and model-predicted power within a given frequency band.

### A small number of eigenmodes capture spatial distributions of alpha and beta band activity

We noticed during our experimentation that only a few eigenmodes appear to contribute substantially to observed MEG alpha and beta patterns. Hence we hypothesized that spatial correlations could be improved by selecting a small subset of eigenmodes. Therefore, we developed a sorting strategy whereby we first rank the eigenmodes in descending order of spatial correlation for a given subject and given frequency band. Then we perform summation over only these eigenmodes according to Eq (10), each time incrementally adding a new eigenmode to the sum. The spatial correlation of these “sorted-summed” eigenmodes against empirical alpha power are plotted in **Figure 5C** as a function of increasing number of eigenmodes; one curve for each subject. The thick black curve represents the average over all subjects. The spatial correlation initially increases as we add more well-fitting eigenmodes, but peaks around, and begins declining thereafter. Addition of the remaining eigenmodes only serves to reduce the spatial correlation. This behavior is observed in almost all subjects we studied.

**Figure 5:**
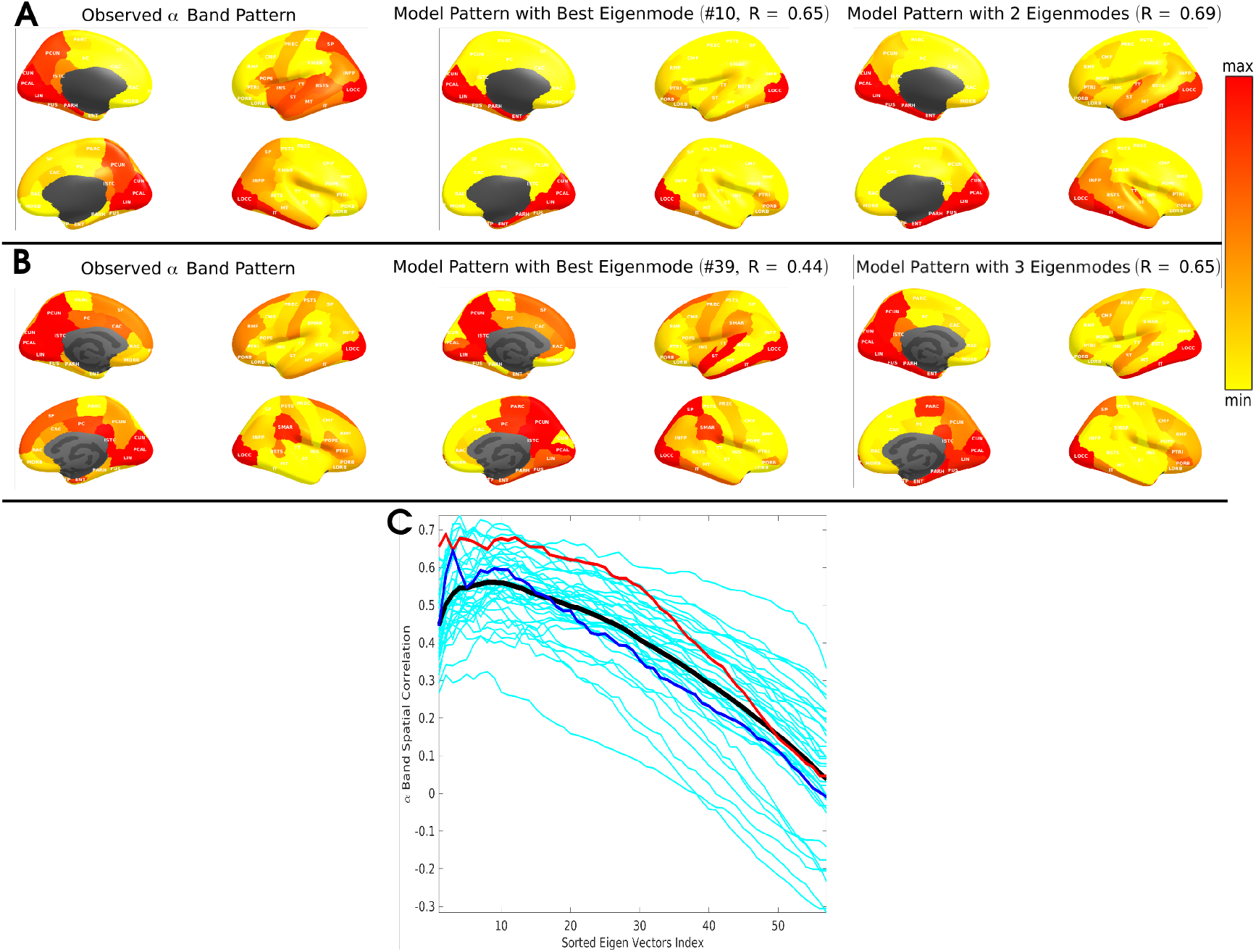
Alpha power spatial distribution depicted by specific spectral graph model eigenmodes. **A & B.** The spatially distributed patterns of alpha band power for two representative subjects are displayed in brain surface renderings. For each four brain panels shown, the medial surface is rendered on the left column while the lateral surface is rendered on the right, the left hemisphere rendering is shown on top while the right hemisphere rendering is shown in the bottom row. **Left column:** The observed MEG spatial distribution pattern for alpha band power showing higher power in posterior regions of the brain. **Middle column:** Spatial distribution of the best matching eigenmode from the spectral graph model. The spatial correlation values are shown on top. **Right column:** Spatial distribution of the best cumulative combination of eigenmodes from the spectral graph model. Spatial correlation values and the number of eigenmodes are shown on top. **C**: Across subject distribution of the alpha band spatial correlation values from spectral graph model simulations for the best fit eigenmodes and the cumulative combination of an increasing number of eigenmodes. Individual subject specific alpha band spatial correlation curves are shown in cyan (n = 36). Panels A and B correspond to the subjects indicated by red and blue curves respectively. Black curve is the average performance across all subjects.

#### Examples of predicted alpha patterns

**Figure 5** shows brain surface renderings of the spatially distributed patterns of alpha band power for two representative subjects. Regions are color coded as a heatmap of regional power scaled by mean power over all regions. The observed MEG spatial distribution pattern of alpha band shows higher power in posterior regions of the brain, as expected, with strong effect size in temporal, occipital and medial posterior areas. This pattern is matched by one of the eigenmodes (#10, shown in middle panel, giving R=0.65), and slightly better by a weighted combination of 2 eigenmodes (R=0.69). However, the model did not reproduce parietal and parieto-occipital components seen in real data. The other subject produced similar results, but with 6 eigenmodes. In this instance, the parietal component seen in real data were reasonably reproduced by the model.

#### Examples of predicted beta patterns

Empirical beta power (**Figure 6**, left) is spread throughout the cortex, especially frontal and premotor cortex. A combination of 4 and 6 best matching eigenmodes produced the best model match to the source localized pattern of two representative subjects, respectively, with R = 0.55 and 0.48. **Figure 6C** shows how the spatial correlation changes as more eigenmodes are used in the “sorted summed” algorithm, analogous to that of alpha pattern. Here too a peak is achieved for a small number of eigenmodes, typically under 10.

**Figure 6:**
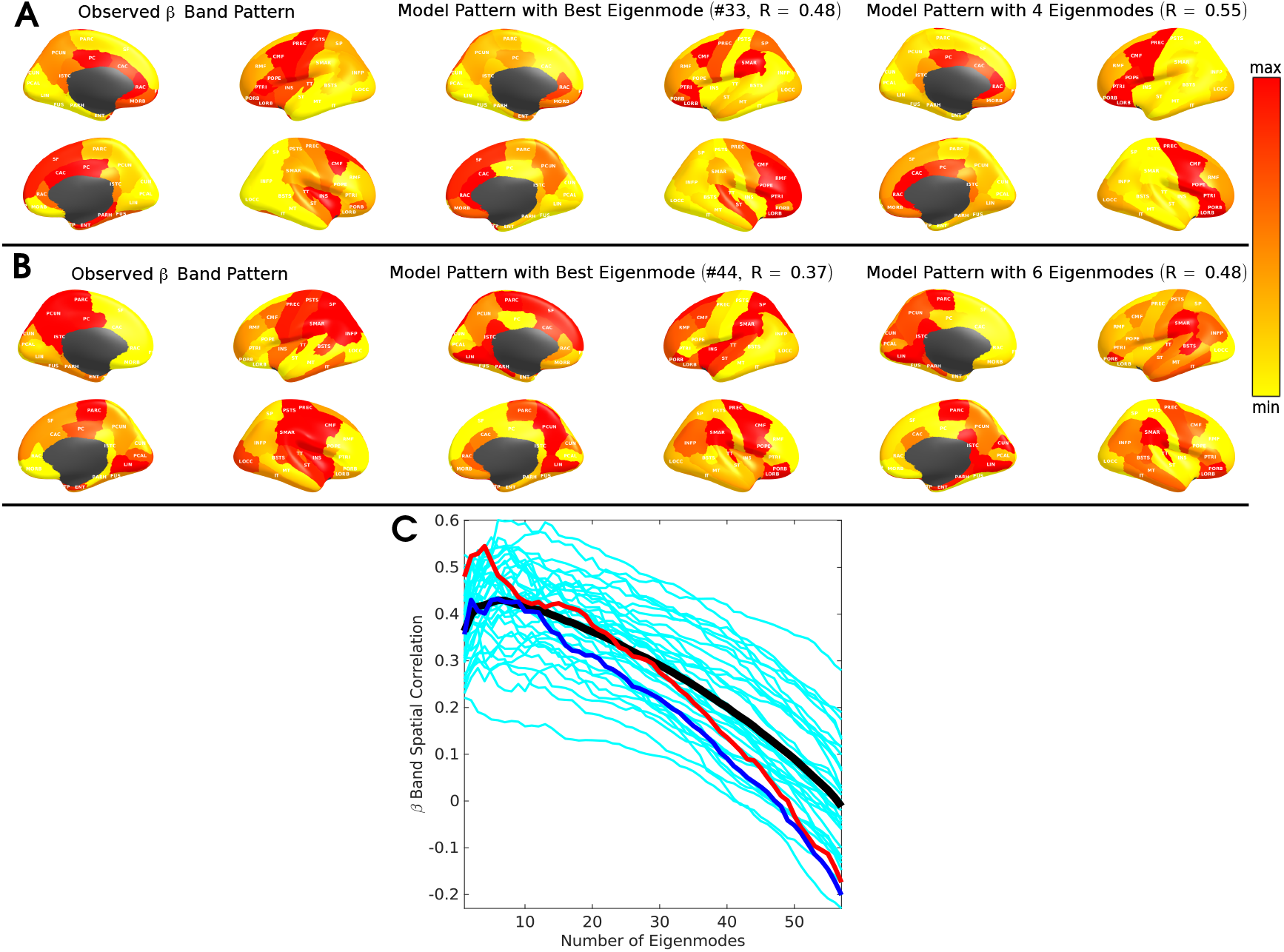
Beta power spatial distributions depicted by specific spectral graph model eigenmodes. Legend is identical to figure 5 but shown for beta power spatial distributions.

### Spatial correlation achieved by the spectral graph model is significantly higher than alternative models

The distribution of peak spatial correlations in the alpha band, using optimized parameters and individual connectomes of all subjects, is plotted in **Figure 7A**. For comparison we show results for four models: a) spectral graph model (SGM) on subject specific individual connectomes (C^Ind^, black); b) SGM on random connectomes with 80% sparsity comparable to individual connectomes (C^Rdm^, blue); c) SGM on geodesic distance based connectomes (C^Dst^, green); and d) a Wilson-Cowan neural mass model (NMM) with subject specific individual connectome (C^Ind^, pink). Analogous results for beta band spatial correlations are contained in **Figure 7B**. Across all subjects the proposed model, SGM on C^Ind^, gives excellent spatial correlations in alpha band (R distribution centered at 0.6) as well as in the beta band (R distribution centered at 0.5). For both alpha and beta spatial distribution patterns, paired t-tests between SGM with C^Ind^ and all other models show that, the SGM with C^Ind^ significantly outperformance all other models, as determined by a paired t-test; p < 0.05 in each case, denoted by asterisk.

**Figure 7:**
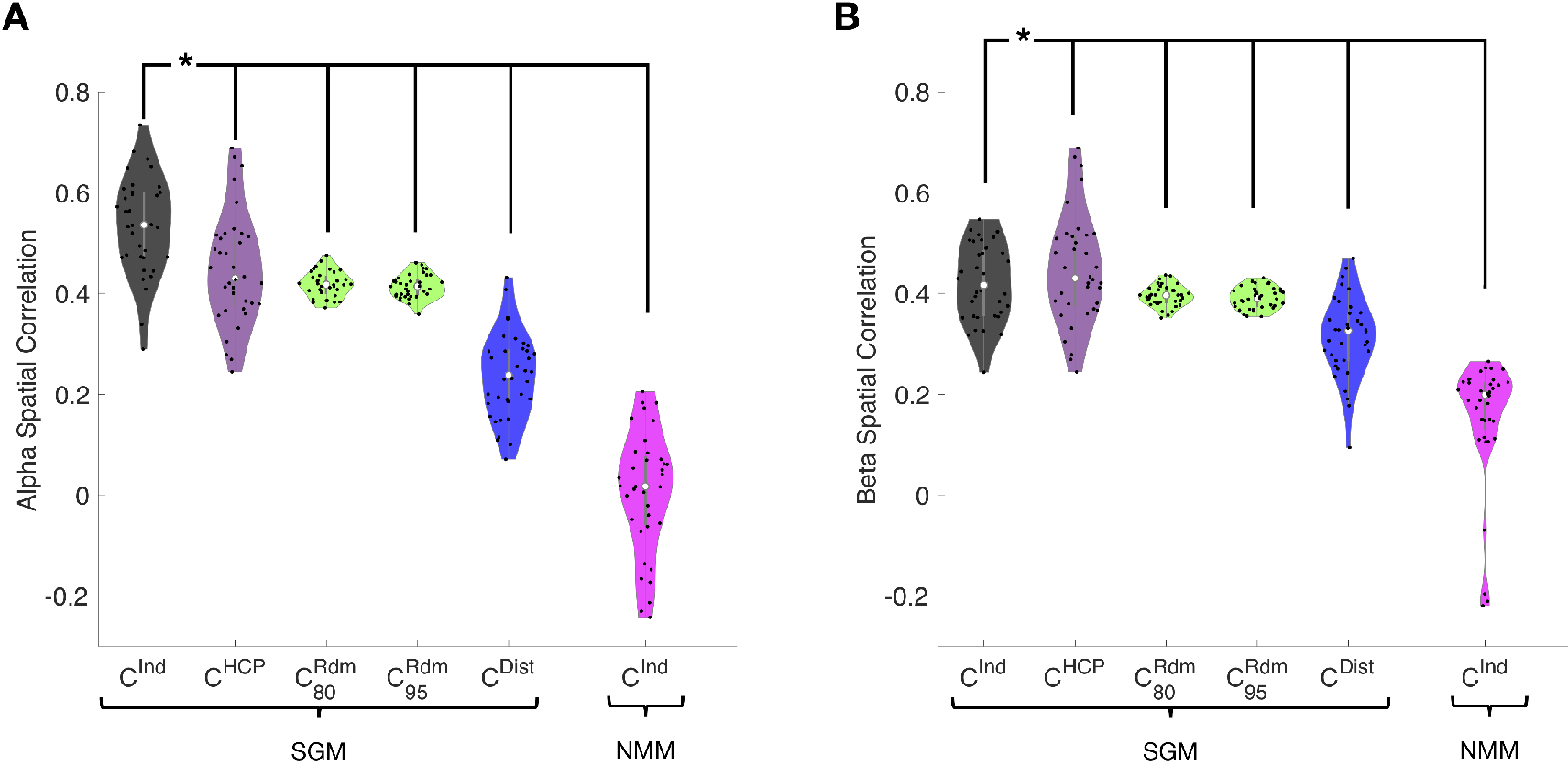
Spatial correlation performance analysis of the spectral graph model. Distribution of the best fit spatial correlations of the spectral graph model across all subjects. **A.** Alpha band spatial correlations. **B.** Beta band spatial correlations. For both panels, spatial correlations are shown for spectral graph model (SGM) with subject specific individual connectomes (C^Ind^, black), random connectomes with 80% sparsity comparable to individual connectomes (C^Rdm^, blue), geodesic distance based connectomes (C^Dst^, green) and for a neural mass model (NMM) with subject specific individual connectome (C^Ind^, pink). For both alpha and beta spatial distribution patterns, paired t-tests between SGM with C^Ind^ and all other models show that, the SGM with C^Ind^ significantly outperformance all other models, as determined by a paired t-test; p < 0.001 in each case, denoted by asterisk.

#### Alternate non-linear model

The Wilson-Cowan neural mass model also did not succeed in predicting the spatial patterns of alpha or beta power, with poor correlations (r centered at 0). This could be because in our implementation we enforced uniform local parameters with no regional variability. However, this is the appropriate comparison, since our proposed model also does not require regionally-varying parameters. Interestingly, the random connectomes and geodesic distance based connectome also appear to have some ability to capture these spatial patterns (r centered at 0.4 and 0.2 respectively), perhaps due to the implicit search for best performing eigenmodes, which on average will give at least a few eigenmodes that look like MEG power purely by chance.

Collectively, we conclude that the graph model is able to fit both the spectral and spatial features of empirical source localized MEG data, and that the optimal fits performed on individual subjects occurs at widely varying subject-specific parameter choices.

## Discussion

The proposed hierarchical graph spectral model of neural oscillatory activity is a step towards understanding the fundamental relationship between network topology and the macroscopic whole-brain dynamics. The objective is not just to model brain activity phenomenologically, but to analytically derive the mesoscopic laws that drive macroscopic dynamics. This model of the structure-function relationship has the following key distinguishing features: *<u>1) Hierarchical</u>:* the model’s complexity depends on the level of hierarchy being modeled: complex, non-linear and chaotic dynamics can be accommodated at the local level, but linear graph model is sufficient at the macro-scale. *<u>2) Graph-based</u>:* Macroscopic dynamics is mainly governed by the connectome, hence linear approximations allow the steady-state frequency response to be specified by the graph Laplacian eigen-decomposition, borrowing heavily from **spectral graph theory**^38–41^. *<u>3) Analytic</u>:* The model is available in closed form, without the need for numerical simulations. *<u>4) Low-dimensional and parsimonious</u>*: Simple, global and universal rules specified with a few parameters, all global and apply at every node, are able to achieve sufficiently complex dynamics. The model is incredibly easy to evaluate, taking no more than a few seconds per brain and to infer model parameters directly from a subject’s MEG data. The optimized model matches observed spectral and spatial patterns in MEG data quite well. No time-consuming simulations of coupled neural masses or chaotic oscillators were needed; indeed, the latter greatly underperformed our model. We report several novel findings with potentially important implications, discussed below.

### Recapitulating regional power spectra at all frequencies

Our main result is the robust demonstration of the model on 36 subjects’ MEG data. The representative examples shown in Figures 3–6 indicate that the graph model recapitulates the observed source localized MEG power spectra for the 68 parcellated brain regions, reproducing the prominent alpha and beta peaks. For each region, the model is also able to predict some characteristics of the full bandwidth power spectra, including what appears to be an inverse power law fall-off over the entire frequency range of interest. However, this aspect will be quantitatively characterized in future work.

We designed a comprehensive parameter optimization algorithm on individual subjects’ MEG data of a suitably defined cost function based on Pearson R statistic as a way to capture all relevant spectral features. Using this fitting procedure, we were able to obtain the range of optimally-fitted parameters across the entire study cohort. As shown in Figure 4A, the range is broad in most cases, implying that there is significant inter-subject variability of model parameters, even if a template connectome is used for all. We tested the possibility that a group-averaged parameter set might also succeed in matching real spectral data on individuals. But as shown in Figures 3B and 4C, this was found to be a poor choice, supporting the key role of individual variability of model parameters (but not variability in the connectome). However, no model is capable of reproducing higher frequencies in the higher beta and gamma range seen in MEG, since by design and by biophysical intuition these frequencies arise from local neural assemblies rather than from modulation by macroscopic networks.

### Revealing sources of heterogeneity in spatial patterns of brain activity

The spatial match between model and data is strongest when the model uses empirical macroscopic connectomes obtained from healthy subjects’ diffusion weighted MRI scans, followed by tractography. The use of “null” connectomes - randomized connectivity matrices of varying levels of sparsity and distance based connectivity matrices, respectively, did far worse than actual human connectomes (Figure 7), supporting the fact that the latter is the key mediator of spatial patterns of real brain activity. The match was also significantly different when using a template HCP connectome versus the individual subject’s own connectomes (Figures 7), and when compared to spatial patterns predicted by an NMM. In conclusion, for the purpose of predicting the spatial topography of brain activity, it is important to use individual connectomes and optimized model parameters.

### Macroscopic brain rhythms are governed by the connectome

A predominant view assumes that different brain rhythms are produced by groups of neurons with similar characteristic frequencies, which might synchronize and act as “pacemakers.” How could this view explain why alpha and beta power are spatially stereotyped across subjects, and why the alpha signal is especially prominent in posterior areas? Although practically any computer model of cortical activity can be tuned, with suitable parameter choice, to oscillate at alpha frequency, e.g.^5,16,20,22,61–63^, none of them are able to parsimoniously recapitulate the posterior origin of alpha. Thus the prominence of posterior alpha might be explained by the hypothesized existence of alpha generators in posterior areas. Indeed, most oscillator models of local dynamics are capable of producing these rhythms at any desired frequency^5,63–66^, and therefore it is common to tweak their parameters to reproduce alpha rhythm. Local networks of simulated multicompartmental neurons can produce oscillations in the range 8-20 Hz^5^, and, in a non-linear continuum theory, peaks at various frequencies in the range 2-16Hz were obtained depending on the parameters^65^. Specifically, the role of thalamus as pacemaker has motivated thalamocortical models^11,16^ that are capable of resonances in various ranges. Neural field models of the thalamocortical loop^16^ can also predict slow-wave and spindle oscillations in sleep, and alpha, beta, and higher-frequency oscillations in the waking state. In these thalamocortical models, the posterior alpha can arise by postulating a differential effect in weights of the posterior versus anterior thalamic projections, e.g.^62^. Ultimately, hypotheses requiring local rhythm generators suffer from lack of parsimony and specificity: a separate pacemaker must be postulated for each spectral peak at just the right location^67^.

An alternative view emerges from our results that macroscopic brain rhythms are governed by the structural connectome. Even with global model parameters, using the exact same local cortical dynamics captured by the local transfer function *H_local_*(*ω*), driven by identically distributed random noise ***P***(*ω*), our model is capable of predicting prominent spectral (Figures 3,4) and spatial (Figures 5,6) patterning that is quite realistic. This is especially true in the lower frequency range: indeed, the model is able to predict not just the frequency spectra in alpha and beta ranges, but also their spatial patterns - i.e. posterior alpha and distributed but roughly frontal beta. Although this is not definitive proof, it raises the intriguing possibility that the macroscopic spatial distribution of the spectra of brain signals *<u>does not require spatial heterogeneity of local signal sources, nor regionally variable parameters</u>*. Rather, it implies that the most prominent *<u>patterning of brain activity (especially alpha) may be governed by the topology of the macroscopic network</u>* rather than by local, regionally-varying drivers. Nevertheless, a deeper exploration is required of the topography of the dominant eigenmodes of our linear model, in order to understand the spatial gradients postulated previously^16,62^.

### Emergence of linearity from chaotic brain dynamics

The non-linear and chaotic dynamics of brain signals may at first appear to preclude deterministic or analytic modeling of any kind. Yet, vast swathes of neuroscientific terrain are surprisingly deterministic, reproducible and conserved across individuals and even species. Brain rhythms generally fall within identical frequency bands and spatial maps^4,16,33^. Based on the hypothesis that the emergent behavior of long-range interactions can be independent of detailed local dynamics of individual neurons^13–18^, and may be largely governed by long-range connectivity^19–22^, we have reported here a minimal linear model of how the brain connectome serves as a spatial-spectral filter that modulates the underlying non-linear signals emanating from local circuits. Nevertheless, we recognize the limitations of a linear model and its inability to capture inherent non-linearities across all levels in the system.

### Relationship to other work

One can view the proposed generative model as a biophysical realization of a dynamic causal model (DCM)^68–72^ for whole brain electrophysiological activity but with very different goals, model dimensionality and inference procedures.

First, the goal of many prior efforts using DCMs is to examine effective connectivity in EEG, LFP and fMRI functional connectivity data, typically for smaller networks^72,73^, or dynamic effective connectivity^74–76^. Hence, they address the second order covariance structures of brain activity. In particular, recent spectral DCM and regression DCM models^77–79^ with local neural masses are formulated in the steady-state frequency-domain, and the resulting whole-brain cross-spectra are evaluated. The goals of these models are to derive model cross-spectra that define the effective connectivity in the frequency domain and are compared with empirical cross-spectra. Based on second-order sufficient statistics, these models attempt to derive effective connectivity from functional connectivity data. These DCMs have so far only been applied to small networks or to BOLD fMRI regime. In contrast, our goal is to examine the role of the eigenmodes of the structural connectome and their influence on power spectral distributions in the full MEG frequency range, and over the entire whole brain. In subsequent work, we intend to extend our efforts to examining effective connectivity but such an effort currently remains outside the scope of the work in this paper. Here, we focus on models that directly estimate the first order effects of observed power spectra and its spatial distributions and compare them with empirical MEG source reconstructions. Our primary motivation is to examine whether spatial distribution of observed power spectra can arise from graph structure of the connectome, hence our focus on the effects of model behavior as a function of the underlying structural connectome - whether it is individualized, template-based, uniform, random or distance based. DCM methods have not reported first order regional power spectra as we do here, nor have they explored how the structural connectome influences model spectral distributions.

Second, our model is more parsimonious compared to most of these above-mentioned models which have many more degrees of freedom because they often allow for regions and their interactions to have different parameters. Our model parameterization, with only a few global parameters, lends itself to efficient computations over fine-scale whole-brain parcellations, whereas most DCMs (with the exception of recent spectral and regression DCMS^77–79^) are suited for examining smaller networks but involve large effective connectivity matrices and region-specific parameters. Furthermore, parameters of our model remain grounded and interpretable in terms of the underlying biophysics, i.e. time constants and conductivities. In contrast, spectral and regression DCM models of cross-spectra have parameters that are abstract and do not have immediate biophysical interpretation.

The third major difference is in the emphasis placed on Variational Bayesian inference in DCM. Since our focus was on exploring model behavior over a small number of global parameters and a set of structural connectomes (whether anatomic or random) of identical sparsity and complexity, it was sufficient to use a *maximum a posteriori* (MAP) estimation procedure for Bayesian inference of our global model parameters with flat non-informative priors with pre-determined ranges based on biophysics. Like most DCM efforts our model can be easily be extended to Variational Empirical Bayesian inference for parameter estimation, for instance to compute a full posterior of the structural connectivity matrix. In such a formulation, we can assume that the observed structural connectome will serve as the prior mean of the connectivity matrix. We reserve such extensions to our future work with this spectral graph model.

### Other limitations and extensions

The model currently examines resting-state activity, but future extensions will include prediction of functional connectivity, task-induced modulations of neural oscillations and causal modeling of external stimuli, e.g. transcranial magnetic and direct current stimulation. The current implementation does not incorporate complex local dynamics, but future work will explore using non-white internal noise and chaotic dynamics for local assemblies. This may allow us to examine higher gamma frequencies. Although our model incorporates latency information derived from path distances, we plan to explore path-specific propagation velocities derived from white matter microstructural metrics such as axon diameter distributions and myelin thickness. Future work will also examine the specific topographic features of the structural connectome that may best describe canonical neural activity spectra. Finally, we plan to examine the ability of the model to predict time-varying structure-function relationships.

### Potential applications

Mathematical encapsulation of the structure-function relationship can potentiate novel approaches for mapping and monitoring brain diseases such as autism, schizophrenia, epilepsy and dementia, since early functional changes are more readily and sensitively measured using fMRI and MEG, compared to structural changes. Because of the complementary sensitivity, temporal and spatial resolutions of diffusion MRI, MEG, EEG and fMRI, combining these modalities may be able to reveal fine spatiotemporal structures of neuronal activity that would otherwise remain undetected if using only one modality. Current efforts at fusing multimodalities are interpretive, phenomenological or statistical, with limited cognizance of underlying neuronal processes. Thus, the ability of the presented model to quantitatively and parsimoniously capture the structure-function relationship may be key to achieving true multi-modality integration.

## Acknowledgements

This work was supported by NIH grants R01EB022717, R01DC013979, R01NS100440, R01DC017696, and UCOP-MRP-17-454755. The template HCP connectome used in the preparation of this work were obtained from the MGH-USC Human Connectome Project (HCP) database (https://ida.loni.usc.edu/login.jsp). The HCP project is supported by the National Institute of Dental and Craniofacial Research (NIDCR), the National Institute of Mental Health (NIMH) and the National Institute of Neurological Disorders and Stroke (NINDS). Collectively, the HCP is the result of efforts of co-investigators from the University of Southern California, Martinos Center at Massachusetts General Hospital (MGH), Washington University, and the University of Minnesota. Additionally, we’d like to acknowledge Pablo F. Damasceno and Megan J. Stanley for contributing to the spectral graph model Python code repository.

## Data Availability Statement

The data and code that support the findings of this study are available from the GitHub repository at https://github.com/Raj-Lab-UCSF/spectrome^80^. The code used to produce basic figures can be run as interactive Jupyter notebooks via Binder^81^. Some raw imaging data, e.g. MRI scans and MEG recordings are not appropriate for public sharing and are too large to be saved in an online repository. However, they could be made available by corresponding author upon reasonable request.

## REFERENCES

1. Hagmann P, Cammoun L, Gigandet X, et al. Mapping the structural core of human cerebral cortex. PLoS Biol. 2008;6(7):1479–1493. doi:10.1371/journal.pbio.0060159

2. Iturria-Medina Y. Anatomical brain networks on the prediction of abnormal brain States. Brain Connect. 2013;3(1):1–21. doi:10.1089/brain.2012.0122

3. Honey CJ, Sporns O, Cammoun L, et al. Predicting human resting-state functional connectivity from structural connectivity. Proc Natl Acad Sci U S A. 2009;106(6):2035–2040. doi:10.1073/pnas.0811168106

4. Freeman WJ, Zhai J. Simulated power spectral density (PSD) of background electrocorticogram (ECoG). Cogn Neurodyn. 2008;3(1):97–103. http://www.pubmedcentral.nih.gov/articlerender.fcgi?artid=2645494&tool=pmcentrez&rendertype=abstract. Accessed September 29, 2015.

5. Liley DT, Alexander DM, Wright JJ, Aldous MD. Alpha rhythm emerges from large-scale networks of realistically coupled multicompartmental model cortical neurons. Network. 1999;10(1):79–92. http://www.ncbi.nlm.nih.gov/pubmed/10372763. Accessed October 5, 2015.

6. Roland PE, Hilgetag CC, Deco G. Tracing evolution of spatio-temporal dynamics of the cerebral cortex: cortico-cortical communication dynamics. Front Syst Neurosci. 2014;8:76. doi:10.3389/fnsys.2014.00076

7. Markounikau V, Igel C, Grinvald A, Jancke D. A dynamic neural field model of mesoscopic cortical activity captured with voltage-sensitive dye imaging. PLoS Comput Biol. 2010;6(9). doi:10.1371/journal.pcbi.1000919

8. Schaffer ES, Ostojic S, Abbott LF. A complex-valued firing-rate model that approximates the dynamics of spiking networks. PLoS Comput Biol. 2013;9(10):e1003301. doi:10.1371/journal.pcbi.1003301

9. Markram H. The blue brain project. Nat Rev Neurosci. 7(2):153–160. doi:10.1038/nrn1848

10. Markram H, Muller E, Ramaswamy S, et al. Reconstruction and Simulation of Neocortical Microcircuitry. Cell. 2015;163(2):456–492. http://www.ncbi.nlm.nih.gov/pubmed/26451489. Accessed October 8, 2015.

11. Izhikevich EM, Edelman GM. Large-scale model of mammalian thalamocortical systems. Proc Natl Acad Sci. 2008;105(9):3593–3598. doi:10.1073/PNAS.0712231105

12. Potjans TC, Diesmann M. The Cell-Type Specific Cortical Microcircuit: Relating Structure and Activity in a Full-Scale Spiking Network Model. Cereb Cortex. 2014;24(3):785–806. doi:10.1093/cercor/bhs358

13. Shimizu H, Haken H. Co-operative dynamics in organelles. J Theor Biol. 1983;104(2):261–273. http://www.ncbi.nlm.nih.gov/pubmed/6227776. Accessed October 5, 2015.

14. Mišić B, Sporns O, McIntosh AR. Communication efficiency and congestion of signal traffic in large-scale brain networks. PLoS Comput Biol. 2014;10(1):e1003427. http://www.pubmedcentral.nih.gov/articlerender.fcgi?artid=3886893&tool=pmcentrez&rendertype=abstract. Accessed October 5, 2015.

15. Mišić B, Betzel RF, Nematzadeh A, et al. Cooperative and Competitive Spreading Dynamics on the Human Connectome. Neuron. 2015;86(6):1518–1529. http://www.ncbi.nlm.nih.gov/pubmed/26087168. Accessed June 18, 2015.

16. Robinson PA, Rennie CJ, Rowe DL, O’Connor SC, Gordon E. Multiscale brain modelling. Philos Trans R Soc Lond B Biol Sci. 2005;360(1457):1043–1050. http://www.pubmedcentral.nih.gov/articlerender.fcgi?artid=1854922&tool=pmcentrez&rendertype=abstract. Accessed September 11, 2015.

17. Destexhe A, Sejnowski TJ. The Wilson-Cowan model, 36 years later. Biol Cybern. 2009;101(1):1–2. doi:10.1007/s00422-009-0328-3

18. Abdelnour F, Voss HU, Raj A. Network diffusion accurately models the relationship between structural and functional brain connectivity networks. Neuroimage. 2014;90:335–347. doi:10.1016/j.neuroimage.2013.12.039

19. Abdelnour F, Raj A, Dayan M, Devinsky OTT. Estimating function from structure in epileptics using graph diffusion model. In: Proceedings of the IEEE International Symposium on Biomedical Imaging.; 2015:Paper ID 528.

20. Nakagawa TT, Woolrich M, Luckhoo H, et al. How delays matter in an oscillatory whole-brain spiking-neuron network model for MEG alpha-rhythms at rest. Neuroimage. 2014;87:383–394. doi:10.1016/j.neuroimage.2013.11.009

21. Jirsa VK, Jantzen KJ, Fuchs A, Kelso JAS. Spatiotemporal forward solution of the EEG and MEG using network modeling. IEEE Trans Med Imaging. 2002;21(5):493–504. doi:10.1109/TMI.2002.1009385

22. Deco G, Senden M, Jirsa V. How anatomy shapes dynamics: a semi-analytical study of the brain at rest by a simple spin model. Front Comput Neurosci. 2012;6:68. doi:10.3389/fncom.2012.00068

23. Achard S, Salvador R, Whitcher B, Suckling J, Bullmore E. A resilient, low-frequency, small-world human brain functional network with highly connected association cortical hubs. J Neurosci. 2006;26(1):63–72. doi:10.1523/JNEUROSCI.3874-05.2006

24. Bullmore E, Bullmore E, Sporns O, Sporns O. Complex brain networks: graph theoretical analysis of structural and functional systems. Nat Rev Neurosci. 2009;10(3):186–198. doi:10.1038/nrn2575

25. Strogatz SH. Exploring complex networks. Nature. 2001;410(6825):268–276. doi:10.1038/35065725

26. van den Heuvel MP, Mandl RCW, Kahn RS, Hulshoff Pol HE. Functionally linked resting-state networks reflect the underlying structural connectivity architecture of the human brain. Hum Brain Mapp. 2009;30(10):3127–3141. doi:10.1002/hbm.20737

27. Hermundstad AM, Bassett DS, Brown KS, et al. Structural foundations of restingstate and task-based functional connectivity in the human brain. Proc Natl Acad Sci. 2013;110(15):6169–6174. doi:10.1073/pnas.1219562110

28. Rubinov M, Sporns O, van Leeuwen C, Breakspear M. Symbiotic relationship between brain structure and dynamics. BMC Neurosci. 2009;10(1):55. doi:10.1186/1471-2202-10-55

29. Ghosh A, Rho Y, McIntosh AR, Kötter R, Jirsa VK. Cortical network dynamics with time delays reveals functional connectivity in the resting brain. Cogn Neurodyn. 2008;2(2):115–120. doi:10.1007/s11571-008-9044-2

30. Abdelnour F, Voss HU, Raj A. Network diffusion accurately models the relationship between structural and functional brain connectivity networks. Neuroimage. 2014;90:335–347. doi:10.1016/j.neuroimage.2013.12.039

31. Abdelnour F, Dayan M, Devinsky O, Thesen T, Raj A. Functional brain connectivity is predictable from anatomic network’s Laplacian eigen-structure. Neuroimage. 2018;172:728–739. doi:10.1016/j.neuroimage.2018.02.016

32. Park H-J, Friston K. Structural and Functional Brain Networks: From Connections to Cognition. Science (80-). 2013;342(6158):1238411–1238411. doi:10.1126/science.1238411

33. He BJ, Zempel JM, Snyder AZ, Raichle ME. The temporal structures and functional significance of scale-free brain activity. Neuron. 2010;66(3):353–369. http://www.pubmedcentral.nih.gov/articlerender.fcgi?artid=2878725&tool=pmcentrez&rendertype=abstract. Accessed July 9, 2015.

34. El Boustani S, Destexhe A. A master equation formalism for macroscopic modeling of asynchronous irregular activity states. Neural Comput. 2009;21(1):46–100. http://www.ncbi.nlm.nih.gov/pubmed/19210171. Accessed June 9, 2015.

35. Wilson HR, Cowan JD. A mathematical theory of the functional dynamics of cortical and thalamic nervous tissue. Kybernetik. 1973;13(2):55–80. http://www.ncbi.nlm.nih.gov/pubmed/4767470. Accessed June 9, 2015.

36. David O, Friston KJ. A neural mass model for MEG/EEG: Coupling and neuronal dynamics. Neuroimage. 2003;20(3):1743–1755. doi:10.1016/j.neuroimage.2003.07.015

37. Xie X, Kuceyeski A, Shah SA, Schiff ND, Nagarajan S, Raj A. Identifiability in connectome based neural mass models. bioRxiv. November 2018:480012. doi:10.1101/480012

38. Larsen R, Nielsen M, Sporring J, Zhang F, Hancock E. Medical Image Computing and Computer-Assisted Intervention – MICCAI 2006. Vol 4191. (Larsen R, Nielsen M, Sporring J, eds.). Berlin, Heidelberg: Springer Berlin Heidelberg; 2006. doi:10.1007/11866763

39. Kondor, Lafferty. Diffusion kernels on graphs and other discrete structures. 2002.

40. Auffarth B. Spectral graph clustering. Univ Barcelona, course Rep. 2007:1–12.

41. A.Ng, M. Jordan YW. On Spectral Clustering: Analysis and an algorithm. Adv Neural Inf Process Syst. 2002:849–856.

42. Galán RF. On How Network Architecture Determines the Dominant Patterns of Spontaneous Neural Activity. Sporns O, ed. PLoS One. 2008;3(5):e2148. doi:10.1371/journal.pone.0002148

43. Goni J, van den Heuvel MP, Avena-Koenigsberger A, et al. Resting-brain functional connectivity predicted by analytic measures of network communication. Proc Natl Acad Sci. 2014;111(2):833–838. doi:10.1073/pnas.1315529111

44. Wang MB, Owen JP, Mukherjee P, Raj A. Brain network eigenmodes provide a robust and compact representation of the structural connectome in health and disease. Jbabdi S, ed. PLOS Comput Biol. 2017;13(6):e1005550. doi:10.1371/journal.pcbi.1005550

45. Abdelnour F, Dayan M, Devinsky O, Thesen T, Raj A. Estimating brain’s functional graph from the structural graph’s Laplacian. In: Papadakis M, Goyal VK, Van De Ville D, eds. SPIE Optical Engineering + Applications. International Society for Optics and Photonics; 2015:95970N. http://proceedings.spiedigitallibrary.org/proceeding.aspx?articleid=2442601. Accessed October 6, 2015.

46. Atasoy S, Donnelly I, Pearson J. Human brain networks function in connectome-specific harmonic waves. Nat Commun. 2016;7:10340. http://www.pubmedcentral.nih.gov/articlerender.fcgi?artid=4735826&tool=pmcentrez&rendertype=abstract. Accessed January 22, 2016.

47. Abdelnour F, Mueller S, Raj A. Relating Cortical Atrophy in Temporal Lobe Epilepsy with Graph Diffusion-Based Network Models. PLoS Comput Biol. 2015;11(10). doi:10.1371/journal.pcbi.1004564

48. Abdelnour F, Raj A, Devinsky O, Thesen T. Network Analysis on Predicting Mean Diffusivity Change at Group Level in Temporal Lobe Epilepsy. Brain Connect. 2016;6(8):607–620. doi:10.1089/brain.2015.0381

49. Raj A, Kuceyeski A, Weiner M. A network diffusion model of disease progression in dementia. Neuron. 2012;73(6):1204–1215.

50. Ferezou I, Haiss F, Gentet LJ, Aronoff R, Weber B, Petersen CCH. Spatiotemporal dynamics of cortical sensorimotor integration in behaving mice. Neuron. 2007;56(5):907–923. doi:10.1016/j.neuron.2007.10.007

51. Polack P-O, Contreras D. Long-range parallel processing and local recurrent activity in the visual cortex of the mouse. J Neurosci. 2012;32(32):11120–11131. doi:10.1523/JNEUROSCI.6304-11.2012

52. Fischl B, Salat DH, Busa E, et al. Whole Brain Segmentation: Automated Labeling of Neuroanatomical Structures in the Human Brain. Neuron. 2002;33:341–355.

53. Owen JP, Li Y-O, Ziv E, et al. The structural connectome of the human brain in agenesis of the corpus callosum. Neuroimage. 2013;70:340–355. http://www.pubmedcentral.nih.gov/articlerender.fcgi?artid=4127170&tool=pmcentrez&rendertype=abstract. Accessed October 20, 2015.

54. Jenkinson M, Beckmann CF, Behrens TEJ, Woolrich MW, Smith SM. FSL. Neuroimage. 2012;62(2):782–790. http://www.ncbi.nlm.nih.gov/pubmed/21979382. Accessed July 9, 2014.

55. Wipf DP, Owen JP, Attias HT, Sekihara K, Nagarajan SS. Robust Bayesian estimation of the location, orientation, and time course of multiple correlated neural sources using MEG. Neuroimage. 2010;49(1):641–655. http://www.pubmedcentral.nih.gov/articlerender.fcgi?artid=4083006&tool=pmcentrez&rendertype=abstract. Accessed October 18, 2015.

56. Zumer JM, Attias HT, Sekihara K, Nagarajan SS. Probabilistic algorithms for MEG/EEG source reconstruction using temporal basis functions learned from data. Neuroimage. 2008;41(3):924–940. http://www.pubmedcentral.nih.gov/articlerender.fcgi?artid=4361188&tool=pmcentrez&rendertype=abstract. Accessed October 18, 2015.

57. Jerbi K, Baillet S, Mosher JC, Nolte G, Garnero L, Leahy RM. Localization of realistic cortical activity in MEG using current multipoles. Neuroimage. 2004;22(2):779–793. doi:10.1016/j.neuroimage.2004.02.010

58. Dalal SS, Zumer JM, Agrawal V, Hild KE, Sekihara K, Nagarajan SS. NUTMEG: a neuromagnetic source reconstruction toolbox. Neurol Clin Neurophysiol. 2004;2004:52. http://www.pubmedcentral.nih.gov/articlerender.fcgi?artid=1360185&tool=pmcentrez&rendertype=abstract. Accessed October 20, 2015.

59. Muldoon SF, Pasqualetti F, Gu S, et al. Stimulation-Based Control of Dynamic Brain Networks. Hilgetag CC, ed. PLOS Comput Biol. 2016;12(9):e1005076. doi:10.1371/journal.pcbi.1005076

60. Kirkpatrick S, Gelatt CD, Vecchi MP. Optimization by simulated annealing. Science. 1983;220(4598):671–680. doi:10.1126/science.220.4598.671

61. Nunez PL, Srinivasan R. A theoretical basis for standing and traveling brain waves measured with human EEG with implications for an integrated consciousness. Clin Neurophysiol. 2006;117(11):2424–2435. http://www.pubmedcentral.nih.gov/articlerender.fcgi?artid=1991284&tool=pmcentrez&rendertype=abstract. Accessed October 5, 2015.

62. Vijayan S, Ching S, Purdon PL, Brown EN, Kopell NJ. Thalamocortical mechanisms for the anteriorization of α rhythms during propofol-induced unconsciousness. J Neurosci. 2013;33(27):11070–11075. http://www.pubmedcentral.nih.gov/articlerender.fcgi?artid=3718379&tool=pmcentrez&rendertype=abstract. Accessed October 5, 2015.

63. David O, Friston KJ. A neural mass model for MEG/EEG: coupling and neuronal dynamics. Neuroimage. 2003;20(3):1743–1755. http://www.ncbi.nlm.nih.gov/pubmed/14642484. Accessed June 11, 2015.

64. van Rotterdam A, Lopes da Silva FH, van den Ende J, Viergever MA, Hermans AJ. A model of the spatial-temporal characteristics of the alpha rhythm. Bull Math Biol. 1982;44(2):283–305. doi:10.1016/S0092-8240(82)80070-0

65. Liley DTJ, Cadusch PJ, Dafilis MP. A spatially continuous mean field theory of electrocortical activity. Netw Comput Neural Syst. 2002;13(1):67–113. doi:10.1080/net.13.1.67.113

66. Spiegler A, Jirsa V. Systematic approximations of neural fields through networks of neural masses in the virtual brain. Neuroimage. 2013;83:704–725. http://www.ncbi.nlm.nih.gov/pubmed/23774395. Accessed October 5, 2015.

67. Nunez PL. A study of origins of the time dependencies of scalp EEG: i--theoretical basis. IEEE Trans Biomed Eng. 1981;28(3):271–280. http://www.ncbi.nlm.nih.gov/pubmed/7228073. Accessed October 5, 2015.

68. Daunizeau J, David O, Stephan KE. Dynamic causal modelling: A critical review of the biophysical and statistical foundations. Neuroimage. 2011;58(2):312–322. doi:10.1016/j.neuroimage.2009.11.062

69. Friston KJ, Preller KH, Mathys C, et al. Dynamic causal modelling revisited. Neuroimage. 2017;(February):0–1. doi:10.1016/j.neuroimage.2017.02.045

70. Pinotsis DA, Hansen E, Friston KJ, Jirsa VK. Anatomical connectivity and the resting state activity of large cortical networks. Neuroimage. 2013;65:127–138. doi:10.1016/j.neuroimage.2012.10.016

71. Razi A, Kahan J, Rees G, Friston KJ. Construct validation of a DCM for resting state fMRI. Neuroimage. 2015;106:1–14. doi:10.1016/j.neuroimage.2014.11.027

72. Pinotsis DA, Geerts JP, Pinto L, et al. Linking canonical microcircuits and neuronal activity: Dynamic causal modelling of laminar recordings. Neuroimage. 2017;146(September 2016):355–366. doi:10.1016/j.neuroimage.2016.11.041

73. Daunizeau J, Kiebel SJ, Friston KJ. Dynamic causal modelling of distributed electromagnetic responses. Neuroimage. 2009;47(2):590–601. doi:10.1016/j.neuroimage.2009.04.062

74. Park HJ, Friston KJ, Pae C, Park B, Razi A. Dynamic effective connectivity in resting state fMRI. Neuroimage. 2018;180(May 2017):594–608. doi:10.1016/j.neuroimage.2017.11.033

75. Van de Steen F, Almgren H, Razi A, Friston K, Marinazzo D. Dynamic causal modelling of fluctuating connectivity in resting-state EEG. Neuroimage. 2019;189(May 2018):476–484. doi:10.1016/j.neuroimage.2019.01.055

76. Preti MG, Bolton TA, Van De Ville D. The dynamic functional connectome: State-of-the-art and perspectives. Neuroimage. 2017;160(December 2016):41–54. doi:10.1016/j.neuroimage.2016.12.061

77. Razi A, Seghier ML, Zhou Y, et al. Large-scale DCMs for resting-state fMRI. Netw Neurosci. 2017;1(3):222–241. doi:10.1162/NETN_a_00015

78. Frässle S, Lomakina EI, Razi A, Friston KJ, Buhmann JM, Stephan KE. Regression DCM for fMRI. Neuroimage. 2017;155(February):406–421. doi:10.1016/j.neuroimage.2017.02.090

79. Frässle S, Lomakina EI, Kasper L, et al. A generative model of whole-brain effective connectivity. Neuroimage. 2018;179(May):505–529. doi:10.1016/j.neuroimage.2018.05.058

80. Xie X, Stanley MJ, Damasceno PF. Raj-Lab-UCSF/spectrome: Spectral Graph Model of Connectomes (Version 0.15). zenodo. November 2019. doi:10.5281/ZENODO.3532497

81. Jupyter P, Bussonnier M, Forde J, et al. Binder 2.0 - Reproducible, interactive, sharable environments for science at scale. In: Akici F, Lippa D, Niederhut D, Pacer M, eds. Proceedings of the 17th Python in Science Conference.; 2018:113–120. doi:10.25080/Majora-4af1f417-011

